# Broadcast, structured, and sequence-dominated: brain-wide computation at cellular resolution in mice

**DOI:** 10.1101/2025.07.30.667641

**Authors:** Michael Schartner, Ari Liu, International Brain Laboratory, Ila Fiete

## Abstract

Until recently, brain activity could be examined either globally, through regional averaging, or locally, at cellular resolution—but not both at once. As a result, regions were characterized as functionally homogeneous units while single neurons were often characterized as heterogeneously and randomly tuned. Here, we overcome this long-standing divide by leveraging the unprecedented density of International Brain Laboratory electrophysiological recordings in mice performing a sensorimotor decision-making task. Computationally integrating these recordings and ordering neurons by functional similarity reveals directly to the eye the full temporal arc of computations unfurling over the phases of a trial and across the brain at cellular and millisecond resolution, providing a dynamical map of the architecture of brain-wide computation. We report that one of the largest shares of brain-wide variance is explained not by task-related sensory responses, integration, or decision related signals, but by neurons encoding internally generated, trial-timed broadcast sequences that carry contextual information and are biased toward hippocampal and entorhinal involvement. We find that individual neurons reliably specialize in interpretable functions, yet neurons with similar functional roles are widely dispersed across regions. Each region contains cells spanning nearly the full repertoire of response types, rendering functional tuning largely unpredictable from cytoarchitectural boundaries or spatial location alone. In particular, even primary visual cortex contains a broad diversity of response types beyond stimulus-locked responses. The dominant discernible organizational distinction is instead between cortical and subcortical processing. Subcortical neurons carry the bulk of internal computations in expert mice, including integration, decision-making, and movement initiation. These patterns support a model of broadcast rather than staged computation, in which signals are widely shared and transformed rather than serially processed. Finally, we find that despite widespread regional mixing, representations at the single neuron level are not randomly mixed. Neurons combine specific features in highly correlated groupings, contradicting the hypothesis of random mixed selectivity at the cellular level. Together, our findings suggest a revised architecture of brain computation: typically interpretable and non-random selectivity at the level of single neurons, coupled to continual brain-wide broadcast rather than staged processing, in which internally structured activity dominates neural dynamics and scaffolds cognition.

## Introduction

From phrenology to modern cytoarchitectural partitioning, the idea that brain areas implement distinct functions – the single area–single function hypothesis – has been a remarkably durable organizing principle. Clear specializations exist across multiple scales: from the sensory periphery, where topographic maps tile visual space(^1^), whisker identity(^2^), and auditory frequency(^3^), to higher-order regions selective for faces^(4;5)^ or semantic content(^6^). At the mesoscale, large-scale recordings and perturbations have further demonstrated differential participation of brain areas across tasks^(7;8;9;10;11;12;13;14)^. Together, these findings have motivated a staged view of computation, in which a region performs a specialized computation on its inputs, passes the result to one or a few downstream areas, and so on — a serial pipeline in which both computation and its products are localized in space, Fig. 1a (left).

**Figure 1.**
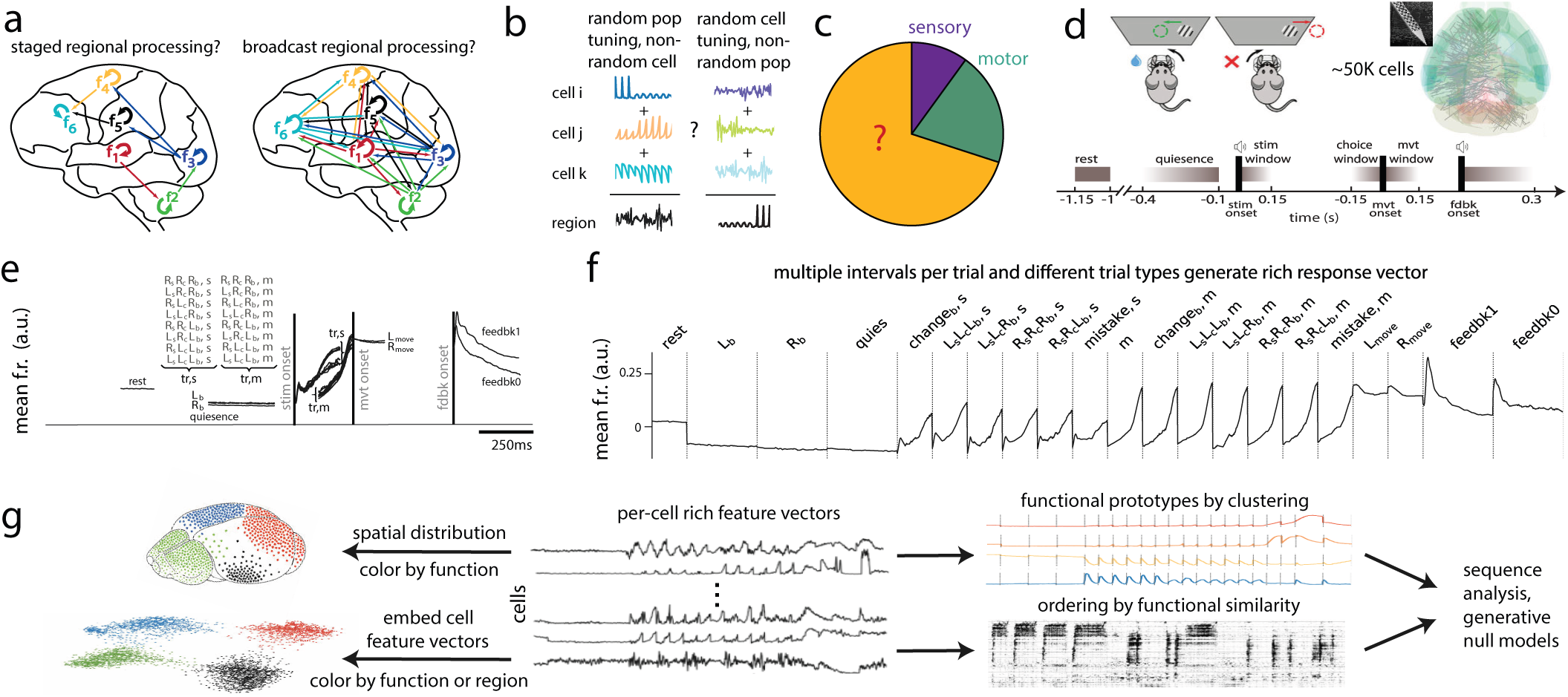
Hypotheses, task structure, and task-aligned rich neural feature vectors across 50K neurons. **a** Schematic of two hypotheses for the architecture of computation in the brain: Left, brain computations are strongly regionalized (spatially modular); or, right, functionally modular computations are performed in distributed multi-regional loops. **b** Schematic of two hypothesis about tuning in the brain: that neurons individually have non-random tuning to specific task features, but at the region level the responses are highly mixed (left) or that individual neurons have highly mixed random tuning to multiple features (mixed selectivity), while possibly combining to yield relatively structured/unmixed region-level function. **c** Up to 9% and 20% of the variance of mouse brain activity during the IBL task is reported to be explained by sensory-decision (stimulus response and integration) and task-related movement (initiation and execution) processes, respectively ((^21^)). The remaining 70% of variance is under-explained. **d** Top left and bottom: schematic of the IBL task and the typical timeline of a trial. A mouse keeps still (quiescence period) to trigger a visual stimulus on the left or right. If it reports the correct stimulus side (*L_e_* or *R_e_*, respectively) by turning a wheel in the appropriate direction, it receives a reward (feedbk 1); otherwise it receives a timeout with white noise over the duration (feedbk 0). After a short rest period, the trial can begin again. Stimulus and feedbk 1 onset are accompanied by an auditory cue. Top right: Neuropixel probe positions across 139 mice, that perform the task at or above expert criterion level, densely tile the left hemisphere. The total yield passing quality control across mice (^21^), used for most of our analyses, is 54719 neurons. In specific analyses that involve cross-validated controls, this number shrinks to 53021. **e** Grand-average brain-wide neural responses aligned to key trial events, separated by trial type (8 types: left, right stim, choice, and block sides) within each event window: within a window, we average the responses of all cells across all trials of the same type (e.g. all feedbk 0 trials within the feedback window, or all trials with left stimulus, left choice, during a right block (*L_s_L_e_R_b_*) in the stimulus window). These curves define peri-event time histograms (PETHs) averaged across all cells. See Tab.1 for details. E.g. in the sensory response and decision windows, stimulus-aligned PETHs are denoted *X_s_X_e_X_b_, s* and movement-aligned PETHs *X_s_X_e_X_b_, m*, where *X* designates *L* or *R*. The stimulus-aligned curves split towards their end in two groups, with the top 4 lines being stimulus side and choice congruent curves while the bottom ones are the 4 incongruent ones, i.e. “mistake” conditions. **f** The grand average *functional feature vector* of all neurons, consisting of 21 PETHs (all windows and all trial types in each window) concatenated then z-scored. **g** Illustration of our analysis pipeline. The per-cell z-scored rich functional feature vectors (middle) are embedded in physical brain space (left, top) or functional space (left bottom), with coloring by clustering on functional similarity or region. We compute the cluster prototypes by averaging feature vectors within a cluster (right, top) to examine functional properties of cells and order cells by similarity (rastermap algorithm (^30^)) to discover temporal patterns (right, bottom). We use the prototypes and rastermap to discover sequences and generate null models for significance testing (far right).

Yetthis view rests substantially on regional averaging and targeted single-neuron sampling with regression of neural responses onto specific task variables. Such approaches have characterized brain areas as functionally homogeneous entities. More recently, recordings that sample several neurons per region have revealed a different picture: in higher cortical areas, neurons can exhibit heterogeneous tuning, often responding to conjunctions of sensory, motor, and contextual variables rather than single features. This observation motivated the mixed selectivity hypothesis, which posits that neurons randomly multiplex features to maximize the dimensionality of population representations and thereby support flexible readout for novel tasks^(15;16;17;18)^. Separately, it has become clear that movement-related signals pervade the brain, explaining more variance in neural activity than sensory-driven or other task-related responses^(19;20;21)^, further challenging the notion of functional segregation. Nevertheless, two critical issues remain unresolved. First, it is unknown how prevalent mixed selectivity is across the brain and whether mixed selectivity is actually random — whether the features a neuron combines are statistically independent — or whether neurons combine features in structured and interpretable ways (Fig. 1b). Second, after accounting for sensory, motor, and decision-related signals, the majority of brain-wide neural variance during a task still remains unexplained (Fig. 1c).

These open questions bear on a fundamental architectural distinction. In the staged model, computation is both local and serial: each region performs a specialized computation on its inputs and passes the result forward, so that the products of a given computation are initially available only to the next stage (Fig. 1a, left). An alternative model preserves local computation but revises both its granularity and its connectivity. In this view, individual brain regions are not unitary computational modules performing a single function. Rather, each region houses multiple local circuits, each participating in distinct computations — and each potentially more deeply interconnected with partner circuits in distant regions than with neighboring circuits within the same anatomical boundary. These distributed, cross-regional circuit partnerships continuously broadcast their computed outputs to the rest of the brain, making them globally available rather than routing them serially to one or a few downstream targets (Fig. 1a, right;^(18;22)^). The broadcast view differs from the staged view in the granularity of the computing units (sub-regional circuits rather than whole regions), the topology of their interactions (distributed loops rather than serial chains), and how the products of computation are shared (globally and in parallel rather than point-to-point). The brain’s massive recurrent connectivity between most regions as well as recent studies highlighting molecular cell-type and connectomic diversity across neurons in a region^(23;24;25;26;27;28)^ are consistent with such an architecture, but until recently the lack of tools for dense, cellular-resolution recordings with broad brain coverage has prevented empirical adjudication.

Here we address these questions using 50,000 Neuropixels-recorded neurons across the mouse brain during a sensorimotor decision-making task in which internal expectations shape perception and guide the progression from stimulus to decision to action^(29;21)^. We define a rich functional feature vector per neuron by concatenating its trial-averaged responses across task phases and trial types, enabling relatively unbiased characterization of each cell’s selectivity. By clustering and sorting this dense brain-wide dataset into an appropriate ordering, it is almost possible to see thoughts writ into neural activity and to trace their progression over time and across the brain. We find that neurons reliably specialize in interpretable functions, yet these functional types are dispersed across the brain rather than localized to specific regions. This dispersion is consistent not with a loss of functional specialization but with a broadcast architecture in which locally computed signals are made globally available. We show that one of the largest shares of neurons belongs to a previously uncharacterized category: internally generated temporal sequences that tile each trial with sparse, contextually modulated activation biased toward hippocampal and entorhinal circuits. Finally, we show that despite regional mixing of functional types, single-neuron representations are not randomly mixed: neurons combine features in structured, correlated groupings, contradicting the random mixed selectivity hypothesis at the cellular level. Together, these findings motivate a reevaluation of many long-held hypotheses about the architecture of computation in the mouse brain.

## Results

On each trial, a visual stimulus appeared on the left or right side of a screen, and mice reported its location by turning a wheel to steer the stimulus to the center (Fig. 1d). Correct responses were rewarded with water; incorrect responses produced a white noise burst and a timeout. Each trial comprised several well-defined phases: a variable-duration rest period, a quiescence requirement (≥200 ms of stillness), stimulus onset, a self-paced decision interval ending at movement onset, and feedback (Fig. 1d). Trials were organized into uncued blocks of 20–100 trials (mean 51), within which the probability of the stimulus appearing on one side was fixed at 0.8 versus 0.2. This probability reversed between successive blocks, creating a latent context that well-trained animals learned to exploit as a prior over stimulus location^(29;31)^. Trials also varied along three binary dimensions — stimulus side (left/right), choice side (left/right), and block side (left/right) — yielding eight trial types that together span the space of sensory, motor, and cognitive variables engaged by the task.

Neural activity was recorded using Neuropixels probes (up to two per mouse) across 139 well-trained animals, yielding ∼50,000 neurons used in our analyses after quality control, with histologically reconstructed positions densely tiling the left hemisphere (Fig. 1d)(^21^). For complete details on the task and dataset, see(^29^) and(^21^), respectively. Despite the variable durations of individual trials, and variable responses of animals and cells, leveraging the trial event structure and trial type structure allows us to obtain standardized and aligned functional responses. To gain a first overview of brain-wide dynamics, we plot the grand average of the activity of all neurons within each task phase, separated by the trial types (Fig. 1e). Several features are apparent. Activity is low during rest and further suppressed during quiescence, but it distinguishes left from right blocks during this pre-stimulus period — reflecting the influence of the learned prior. Stimulus onset produces a brief transient (∼20 ms latency) of modest amplitude, followed by a much larger ramping signal lasting 250–300 ms. The ramp magnitude is modulated by trial type, specifically on whether the stimulus side is concordant or discordant with the block side. When aligned to movement onset, these ramps nearly converge across trial types, consistent with an accumulation-to-bound process, though with a possible small difference in the bound for concordant and discordant trials. The ramp plateaus at a sustained level during motor execution. The largest overall responses in the brain occur at feedback onset, for both positive and negative outcomes.

To move beyond population averages and characterize the functional identity of individual neurons, we constructed a rich feature vector for each cell. This vector concatenates the neuron’s average firing rate time course (PETH) under 21 conditions spanning all task phases and trial types; we illustrate this rich feature vector for the grand average response from Fig. 1e (Fig. 1f). The rich feature vector covers: rest and quiescence periods; pre-stimulus block-side responses; stimulus-aligned responses for each of the four correct trial types, for block-change trials, and for mistake trials; movement-aligned responses for the same conditions; left and right movement execution; and positive and negative feedback (see Table 1 for complete definitions). Though it is possible in principle to further subdivide conditions (e.g. by the type of mistake trial: mistakes right after a block change or deep into a block; or mistakes on left versus right blocks, etc.), we balanced choosing as large a set of conditions while keeping as many sessions as possible (parsing conditions more finely leads to the rejection of more sessions because the number of trials per condition drops below our inclusion threshold). The aligned feature vectors for all neurons across animals constitutes a single “supersession”, whose average across all neurons (Fig. 1f) is equivalent to concatenating the curves in Fig. 1e and then z-scoring.

**Table 1.** PETH definitions. Each PETH is defined by an alignment event, the trials to average, and the window. The two times in the window are relative to the alignment event, with positive values if after the event and negative values if before the event. “Block change” indicates trials within 3 trials of a block switch. “Deep block” indicates averaged trials must be at least 3 trials after a block change.

| Name | Alignment Event | Trial Types to Average | Window (s) |
| --- | --- | --- | --- |
| rest | stimOn_times | All trials | -1.15 to -1 |
| L<sub>b</sub> | stimOn_times | ProbabilityLeft = 0.8 (block left) | -0.4 to -0.1 |
| R<sub>b</sub> | stimOn_times | ProbabilityLeft = 0.2 (block right) | -0.4 to -0.1 |
| quies | stimOn_times | All trials | -0.4 to -0.1 |
| change<sub>b, s</sub> | stimOn_times | All trials within 3 trials of a block change | 0 to 0.15 |
| L<sub>s</sub>L<sub>c</sub>L<sub>b, s</sub> | stimOn_times | Stim left, choice left, block left, deep block | 0 to 0.15 |
| L<sub>s</sub>L<sub>c</sub>R<sub>b, s</sub> | stimOn_times | Stim left, choice left, block right, deep block | 0 to 0.15 |
| R<sub>s</sub>R<sub>c</sub>R<sub>b, s</sub> | stimOn_times | Stim right, choice right, block right, deep block | 0 to 0.15 |
| R<sub>s</sub>R<sub>c</sub>L<sub>b, s</sub> | stimOn_times | Stim right, choice right, block left, deep block | 0 to 0.15 |
| mistake, s | stimOn_times | All 4 mistake types | 0 to 0.15 |
| m | firstMovement_times | All trials | -0.15 to 0 |
| change<sub>b, m</sub> | firstMovement_times | All trials within 3 trials of a block change | -0.15 to 0 |
| L<sub>s</sub>L<sub>c</sub>L<sub>b, m</sub> | firstMovement_times | Stim left, choice left, block left | -0.15 to 0 |
| L<sub>s</sub>L<sub>c</sub>R<sub>b, m</sub> | firstMovement_times | Stim left, choice left, block right | -0.15 to 0 |
| R<sub>s</sub>R<sub>c</sub>R<sub>b, m</sub> | firstMovement_times | Stim right, choice right, block right | -0.15 to 0 |
| R<sub>s</sub>R<sub>c</sub>L<sub>b, m</sub> | firstMovement_times | Stim right, choice right, block left | -0.15 to 0 |
| mistake, m | firstMovement_times | All 4 mistake types | -0.15 to 0 |
| L<sub>move</sub> | firstMovement_times | Choice left | 0 to 0.15 |
| R<sub>move</sub> | firstMovement_times | Choice right | 0 to 0.15 |
| feedbk1 | feedback_times | Positive feedback | 0 to 0.3 |
| feedbk0 | feedback_times | Negative feedback | 0 to 0.3 |

**Table 2.** Mapping of Functional Networks to Selected Literature, Mouse Sample Sizes, and Experimental States.

| Network Domain | Citations | Sample Size (*N*) | Experimental Condition / State |
| --- | --- | --- | --- |
| **Default Mode** | (10) | *N* = 18 | Anesthetized (isoflurane + medetomidine), resting-state |
|  | (42) | *N* = 610 | Varied (awake and diverse anesthesia protocols across 14 sites) |
|  | (41) | *N* = 20 | Anesthetized (medetomidine), resting-state |
|  | (48) | *N* = 45 | Anesthetized (isoflurane + medetomidine), resting-state |
|  | (45) | *N* = 19 | Anesthetized (isoflurane + pancuronium bromide), resting-state |
| **Limbic / Hippocampal** | (43) | *N* = 15 / 15 | Awake vs. Anesthetized (halothane), resting-state |
|  | (46) | *N* = 15 | Anesthetized (halothane), resting-state |
|  | (42) | *N* = 610 | Varied (awake and diverse anesthesia protocols across 14 sites) |
|  | (47) | *N* = 50 | Anesthetized (5 groups: isoflurane, propofol, medetomidine, combinations) |
| **Salience** | (40) | *N* = 34 | Awake, resting-state |
|  | (42) | *N* = 610 | Varied (awake and diverse anesthesia protocols across 14 sites) |
|  | (45) | *N* = 19 | Anesthetized (isoflurane + pancuronium bromide), resting-state |
| **Sensorimotor** | (44) | *N* = 9 | Awake, resting-state |
|  | (46) | *N* = 15 | Anesthetized (halothane), resting-state |
|  | (41) | *N* = 20 | Anesthetized (medetomidine), resting-state |
|  | (42) | *N* = 610 | Varied (awake and diverse anesthesia protocols across 14 sites) |
| **Associated Cortical / Laterocortical** | (41) | *N* = 20 | Anesthetized (medetomidine), resting-state |
|  | (48) | *N* = 45 | Anesthetized (isoflurane + medetomidine), resting-state |
|  | (42) | *N* = 610 | Varied (awake and diverse anesthesia protocols across 14 sites) |
|  | (45) | *N* = 19 | Anesthetized (isoflurane + pancuronium bromide), resting-state |

In what follows, we analyze the structure of the full supersession matrix of the feature vectors of ∼50,000 neurons through clustering, sorting, manifold embedding, and associated quantifications (Fig. 1g).

### Unsupervised discovery of structured and interpretable rich neural response categories

Do the ∼50,000 constructed feature vectors tile a continuous, unstructured response space, or do neurons exhibit discrete functional groupings? To address this, we applied k-means clustering (*N* = 25 clusters) to the full supersession matrix (*N_c_* = 54,719 neurons × *N_T_* bins, where *N_T_* = 1872 (with *t_bin_* = 125*ms*) is the length of each feature vector), obtaining 25 prototype response vectors — each the average feature vector of its member neurons (Fig. 2a).

**Figure 2.**
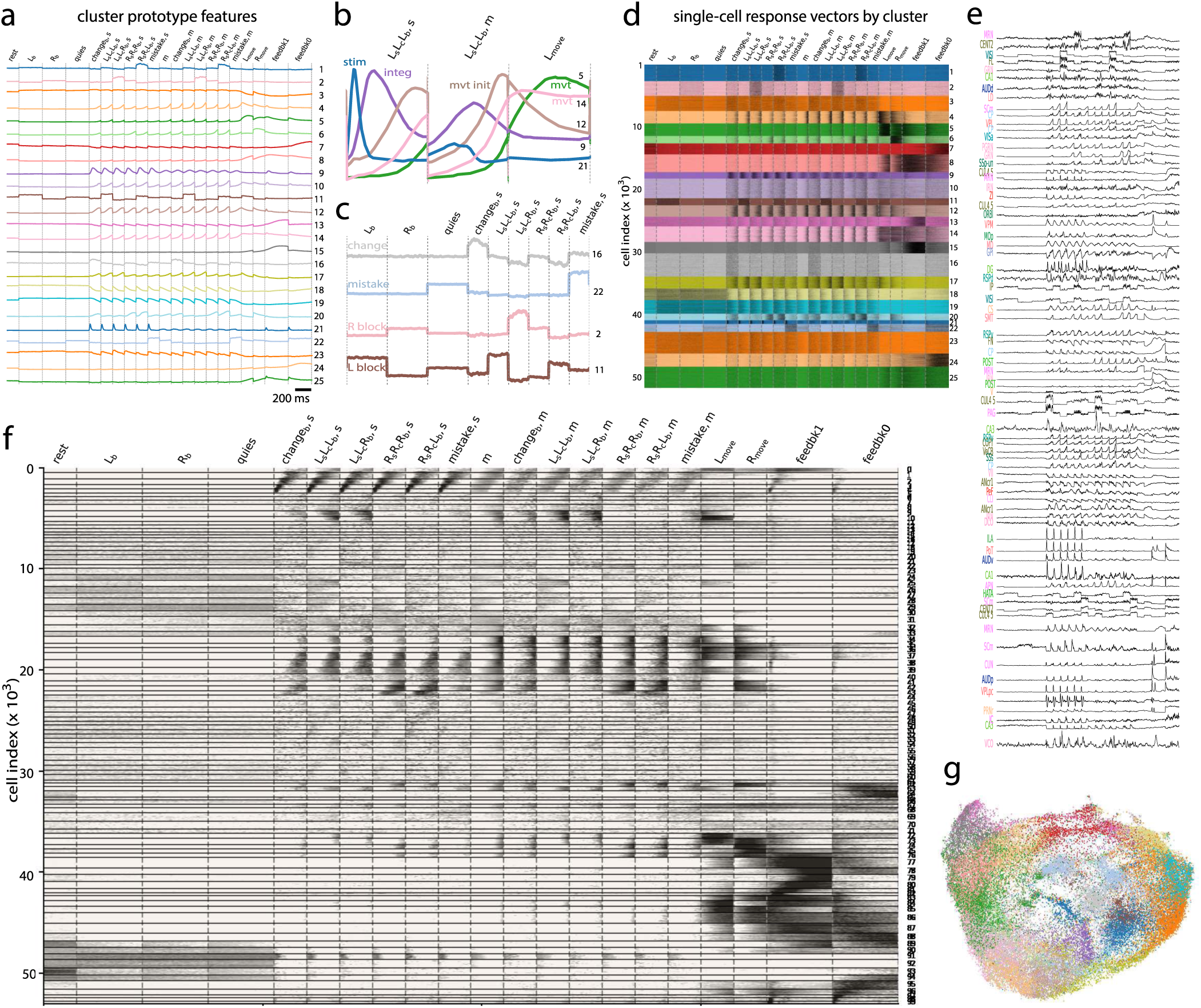
Brain-wide activation across trial types and trial intervals reveals functionally interpretable features at the population and single-neuron levels. a Average response vectors of 25 k-means clusters, note e.g. the stimulus responsive cluster number 21 and the positive feedback responsive clusters 13 and 15. b Zoomed-in view of the average response vectors of cluster 21, 9, 12, 14, and 5, showing response patterns of a stimulus response cluster, a ramping/integration cluster, a movement initiation cluster, and two movement clusters, respectively. c Zoomed-in view of the average response vectors of cluster 16, 22, 2, and 11, encoding response for surprise/block change, mistake, right block, and left block trials, respectively. d Single cell response raster plot of all neurons, sorted and colored by k-means clusters as in a. e Example single cell feature vectors from different Beryl regions (labels on the left, see Methods and Fig. S12 for anatomical parcellation information) across the whole brain, showing interpretable functional responses similar to the cluster-level averages in a. f Feature vectors of all ∼ 50 k cells sorted by similarity via rastermap (^30^) into 100 clusters, revealing great diversity in activity and clear patterns. Black is high firing rate and white is low. Black horizontal lines indicate cluster boundaries, and the clusters are numbered on the right. g UMAP embedding into a two-dimensional space of all ∼ 50 k cells’ feature vectors, colored by k-means cluster identity as in a.

Prior analyses of IBL and similar decision tasks have typically classified neurons into a small set of hand-defined categories: stimulus-selective, choice-selective, and feedback-selective neurons(^21^). Our unsupervised prototypes recover these, but decompose them more finely and reveal additional structure and features. We assigned prototypes to task events and functional roles by identifying the trial phase(s) and alignments for which each prototype’s response was most sharply peaked. For instance, the blue and purple curves (Fig. 2b) exhibit elevated firing in both the stimulus onset and pre-movement periods, but their response peaks are more temporally defined (narrowly peaked) and higher in amplitude when aligned to stimulus onset, identifying them as stimulus-related responses. By contrast, the beige and pink curves (Fig. 2b) are also active in both periods but peak far more sharply when movement-aligned, identifying them as movement-associated responses. Different movement-related categories (e.g. movement choice, preparation or initiation versus movement generation) can be distinguished based on the timing of the response peak prior to movement and whether activity decays going into the actual movement period or continues or increases. We observe a transient stimulus-response cluster (#21) that is active not only at stimulus onset at the beginning of each trial but also at the start of the feedback intervals where an auditory tone is emitted; a ramping cluster consistent with evidence integration (#9); a movement-initiation cluster (#12); and two distinct movement-execution clusters (#5, #14) (Fig. 2b, top), the latter of which exhibits a combined movement-and-feedback response.

Beyond these sensorimotor categories, we found several prototypes that appear to encode internal or cognitive states (Fig. 2c). These include: a block-change cluster, in which during-stimulus activity is elevated on trials shortly following a block switch, consistent with a surprise or prediction-error signal; a mistake-encoding cluster for mistake trials that exhibits elevated quiescence-period activity; left-and right-block clusters encoding the latent prior during quiescence; positive and negative feedback clusters (#15 and #24, respectively, Fig. 2a); and two conjunctive movement-and-feedback clusters (#8, #14). (Extended Data Fig. 9 in(^31^) demonstrates that movement confounds are manageable for that analysis).

A natural concern with clustering is that it can impose discrete boundaries on a continuous distribution with prototypes that are the per-tile averages of a tiling of the continuous space, rather than reflections of intrinsic structure. We evaluated this in several ways. First, sorting neurons by cluster membership and inspecting the full per-neuron supersession raster (Fig. 2d) reveals that individual cells within each cluster exhibit response patterns that clearly resemble their assigned prototype — the prototypes are not mere averages over heterogeneous responses. Second, examining traces from individual neurons within each cluster (Fig. 2e) confirms that single cells show distinct peaks and response profiles matching the prototypes, consistent with actual functional specialization. Thus, though there is a diversity of neural responses within each functional cluster, the clusters nevertheless are meaningful partitions of single neurons, and single neurons exhibit specific structured response patterns with functional meaning relative to the task, in coherent multi-neuron groupings. Third, a nonlinear embedding of the feature vectors (UMAP; Fig. 2g) reveals groupings in the low-dimensional embedding space which align with k-means cluster assignments (colors), indicating that the discrete k-means clusters reflect geometric structure in the high-dimensional response space rather than arbitrary partitioning. Varying the number of clusters (*N* = 10 to 100) yields finer-grained categories but does not qualitatively alter this picture (Fig. S1).

In sum, neurons across the brain can be clustered into structured, interpretable, and rich functional categories that extend beyond previously defined response types.

### Cell ordering reveals brain-wide computational dynamics over the arc of decision-making at cellular resolution

The k-means analysis above assigned each neuron to one of 25 discrete clusters. While this reveals interpretable functional categories, it collapses within-cluster heterogeneity and obscures finer-grained temporal structure — for instance, continuous variation in response latency among neurons with a shared functional role. To obtain a higher-resolution and temporally ordered view, we organized the ∼50,000 neurons according to rastermap(^30^) using 100 functonal clusters, which orders clusters based on their similarity.

The result is a single image of brain-wide computation unfolding at cellular resolution across trial phases, trial types, and time (Fig. 2f). The rastermap recapitulates the discrete functional categories found by k-means — for instance, feedback-preferring neurons form a large coherent block (clusters 76–90, lower right), and movement neurons form another (clusters 34–42). But it also reveals structure that clustering alone cannot capture. Within many functional blocks, responses exhibit continuous and systematic temporal variation across neurons. For example, one population encodes a rapid stimulus-onset transient (dark portion of the whorl near cell index ∼2,500), grading smoothly into a second population that extends the stimulus response at longer latencies (upper portion of the whorl, cells 1–2,500). This continuous temporal tiling shows that neurons inhabit a spectrum of roles, with preserved temporal structure or relative latencies across trial types.

Beyond confirming and refining the discrete categories, the rastermap reveals a previously uncharacterized response type: large sets of neurons exhibiting faint diagonal bands spanning the interval from trial onset to movement onset (clusters 5–7, 11–20, 45–54, and 56–59). These diagonal patterns — suggestive of sequential activation in which successive neurons fire at successive time points — are strikingly consistent across trial types, appearing in multiple columns of the rastermap. We examine these sequences in detail in the next section.

### Reliable, brain-wide contextual sequences scaffold experience during each trial

When neurons are sorted by functional similarity using rastermap, a striking feature emerges: large blocks of neurons (clusters 5–7, 11–20, 45–54, 56–59) appear to exhibit sequential activation patterns — diagonal bands in the rastermap — spanning the interval from trial onset to movement onset (Fig. 2f). Here we examine and characterize these sequences in detail.

We performed an explicit cross-validation by splitting all trials into two halves. We applied rastermap to one half (“train”; Fig. 3a, left), identifying strong diagonal bands. We then applied the neuron ordering learned from the train half to the held-out test half (see Fig. S2 for full train/test rastermaps). The sequences persist clearly in this fully disjoint and independent trial set (Fig. 3a, middle), and survive averaging across trial types (Fig. 3a, right). In fact, each non-mistake trial type within the rich feature vector is constructed from entirely different non-overlapping trials: thus, sequential structure across trial types even within a set (within train or within test halves) constitutes an intrinsic cross-validation. Though neurons can always be sorted to create sequence-like patterns within a trial type, cross-trial type sequences cannot result as an artifact of such neural ordering. Given this fact, we can also return to the train set to interpret some of its features. Within the train set, the sorting process identifies three types of sequences (three diagonal bands within each trial type), with the diagonal bands most prominent in trial types involving some form of surprise — discordant trials (stimulus side opposite to block side, i.e. *L_s_L_e_R_b_, s* and *R_s_R_e_L_b_, s*) and block-change trials — but persisting across all remaining trial types; mistake conditions have the weakest bands, but the diagonal band in block-change trials appears relatively strong in mistake trials, indicating mistakes are closely related to surprise upon block-change. Despite these differences, averaging across trial types within the test set confirms the existence of a core sequence backbone common to all conditions (Fig. 3a, right).

**Figure 3.**
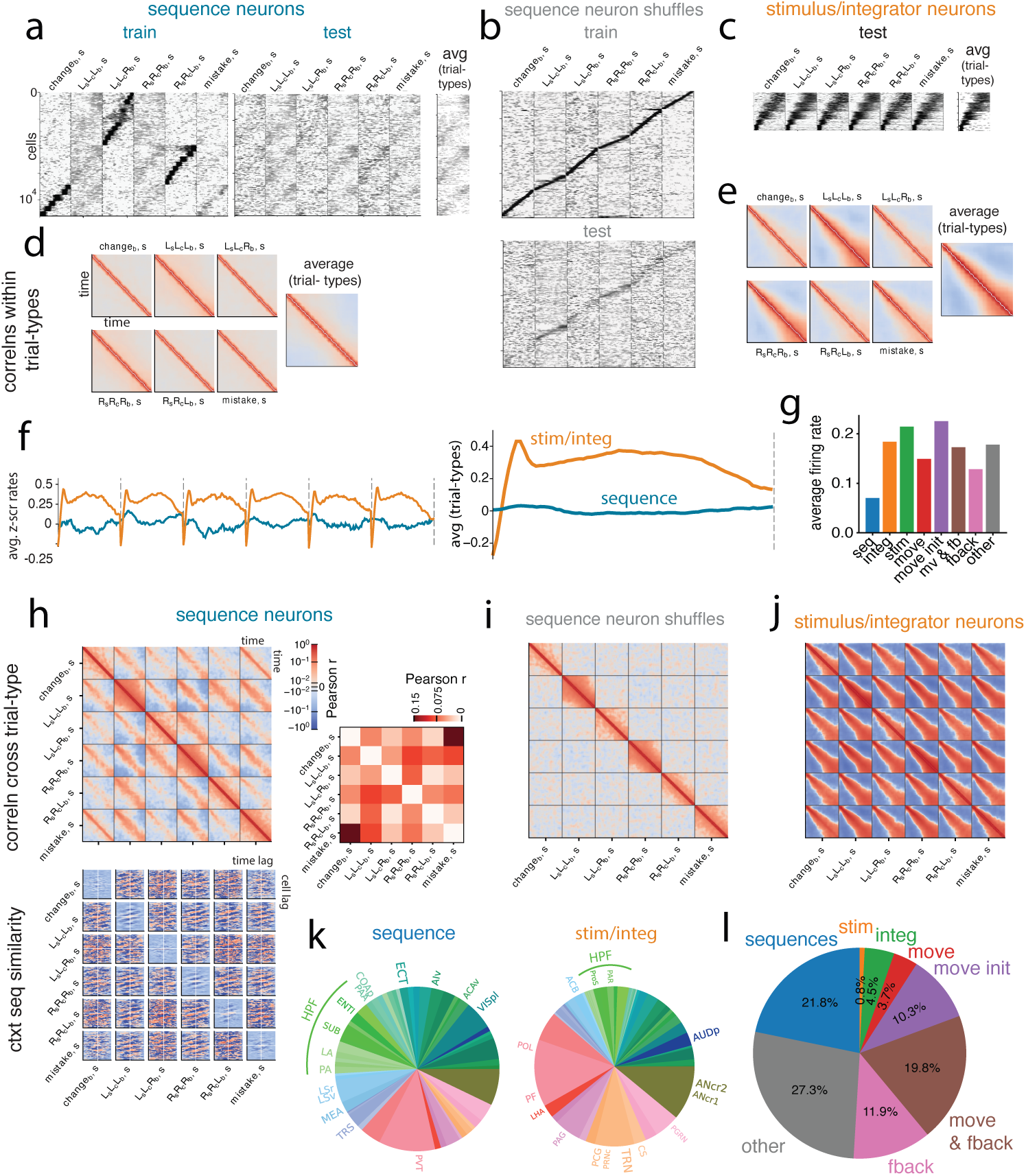
Internally generated brain-wide contextual neural sequences. **a** Sequence neurons identified and ordered by the rastermap algorithm and verified by cross-validation (Fig. S2 shows the full train/test rastermaps). Left: raster plot based on the train set (odd-numbered trials), showing three groups of strong contextual specific sequence neurons for block change (change_b_, s), L_s_L_c_R_b_, s, and R_s_R_c_L_b_, s trials, and the same neurons also show sequential responses across all stimulus-aligned trial conditions. Right: the test set (even-numbered trials) raster plot for stimulus-aligned trial conditions and average over these conditions. The strong contextual responses did not survive cross-validation, but the sequences across all trial conditions are still evident in the test set. **b** Sequence neurons with independently shuffled row ordering within each trial type, and re-sorted using rastermap with cross-validation. Top: the train set of trials which rastermap is run on after row-shuffling; bottom: test set sorted by the resulted ordering of train set, showing faint sequences within each trial type, but they do not generalize across trial types as in **a** (column-shuffling prior to rastermap does not result in sequences, see Fig. S4a). **c** Stimulus response and integrator neurons, identified and ordered by rastermap, and cross-validated (only the test set shown here). We show their raster plot for all stimulus-aligned trial conditions as well as the average across these trial types. **d** Time-time correlation matrices of sequence neuron PETHs within each of the trial types, and of the average PETHs across the different trial types have narrow and constant bands at the diagonals (exact diagonals set to 0 for visualization). **e** Time-time correlation matrices of stimulus response and integrator neuron PETHs within each of the trial types, and of the average PETHs across the different trial types, showing variable widths around the diagonals, unlike for sequence cells; diagonal values set to 0 for visualization. **f** Average z-scored PETHs of the sequence neurons (blue) and stimulus response + integrator neurons (orange), for each individual trial type (left) and averaged over trial types (right). **g** Average firing rate of functional groups defined by rastermap algorithm (see Methods for definition), across all concatenated trial windows in the full feature vectors. **h** Correlation structure across trial types in the sequence neurons. Top left: time-time correlation matrix of sequence neuron PETHs across all stimulus-aligned trial types concatenated together. Colors indicate the Pearson correlation score and are shown in log-scale. Top right: overall Pearson correlation of sequence neurons across trial conditions, constructed by taking the cell-by-time matrices of each trial condition and calculating pair-wise correlation among them. The diagonal values are set to 0 for visualization. Bottom: two-dimensional cross-correlograms of sequence neurons across trial conditions. Each cross-correlogram is constructed by taking a pair of trial conditions’ cell-by-time matrices, and computing the correlation at each time lag and cell lag. Colors indicate correlation strength and are shown in log-scale. **i** Time-time correlation matrix of row-shuffled sequence neuron PETHs, in the test set of trials as in **b**, across all stimulus-aligned trial types concatenated together. The same analysis on the train set is qualitatively the same and thus not shown. **j** Time-time correlation matrix of stimulus response and integrator neuron PETHs across all stimulus-aligned trial types concatenated together. Colors shown in log-scale. **k** Regional composition of sequence neurons (left) and stimulus response and integrator neurons (right), colored by Beryl regions. The top 15 regions are annotated with their acronyms, and hippocampal regions are indicated as’HPF’. **l** Functional composition of the 50k cells included in analysis, as identified by rastermap algorithm.

Randomly shuffling time indices (columns) independently within each trial type before applying rastermap abolishes all visible sequential structure (Fig. S4a). Shuffling neural indices separately per trial type (maintaining their temporal responses within trial type), then splitting the data into a train and test set and sorting the resulting test set with rastermap can indeed induce sequences within individual trial types (Fig. 3b), but — unlike the real data — this ordering does not generalize across trial types (Fig. 3a), confirming that the cross-trial-type consistency of the sequences depends on stable, neuron-specific temporal tuning.

The sequence neurons are distinct from other functional groups identified by rastermap. Stimulus-responsive and integrator neurons occupy separate clusters (clusters 1–4 and 38; Fig. 3c), as do movement-initiation neurons (clusters 10, 32, 34–37, 39, 41–42, and 61, as numbered in Fig. 2f right y axis; see Methods and Fig. S3 for definition of all rastermap-defined functional types). All subsequent analyses use only held-out test trials with the training-derived ordering of sequence neurons.

The distinction is confirmed by the temporal autocorrelation structure. For sequence neurons, the autocorrelation matrix within each trial type exhibits a band of constant width along the diagonal (Fig. 3d) — the signature of a population tiling time or the represented abstract space uniformly. This is reminiscent of hippocampal place cells and time cells, which tile space or delay intervals with approximately uniform fields and population density^(32;33;34)^. By contrast, stimulus-integrator neurons show a broadening autocorrelation profile along the diagonal (Fig. 3e), reflecting temporal dispersion — an initially sharp stimulus-locked response that progressively smears as integration unfolds. Column-shuffled controls lack any temporal structure (Fig. S4b).

Two additional distinctions appear in functional neuron group-averaged population activities over time. Stimulus and integrator neurons exhibit a transient onset peak followed by a growing ramp that decays — the expected signature of stimulus detection followed by evidence accumulation. Sequence neurons, by contrast, maintain a nearly constant total firing rate across the trial (Fig. 3f). Additionally, the mean firing rate of sequence neurons is substantially lower than that of other functional response categories including groups that seem to encode stimulus, integrator, movement, and feedback neurons (Fig. 3g; see Methods and Fig. S3 for examples of these rastermap-defined functional groups). The flat population rate profile, combined with individual neurons that each fire sparsely at specific trial times, is what one would expect from neurons encoding abstract sequences unrelated to specific stimuli. It implies that the sequence population acts as an abstract temporal basis set: individual neurons tile time while their sum remains approximately constant. These properties are reminiscent of the tiling of time by HVC premotor sequence cells(^35^) or of space or time by hippocampal place or time cells, respectively^(32;36)^.

Besides carrying shared information across trial types (Fig. 3a, right), the sequences also contain trial-type-specific information. Cross-correlations computed across trial types reveal a consistent diagonal structure of finite width in all trial-type pairs (Fig. 3h, top-left), confirming the shared backbone. However, the strength of cross-correlation varies systematically with trial-type similarity. The weakest cross-correlation occurs between the two opposite discordant trials, R_s_R_c_L_b_ and L_s_L_c_R_b_, while mistake sequences correlate most strongly with block-change sequences (Fig. 3h, top-right). This pattern suggests a hierarchical contextual encoding: the sequences first distinguish surprise from non-surprise conditions, then further differentiate specific trial types within each category.

To characterize the full spatiotemporal structure, we computed two-dimensional cross-correlograms between all pairs of trial types, treating each trial-type’s rastermap submatrix (neurons × time bins) as a two-dimensional image as commonly done in the hippocampal field (Fig. 3h, bottom; see Methods for details). Across all condition pairs, the correlograms exhibit a consistent slope and a 7-band structure, corresponding to three repeated sequence motifs in the rastermap that align with the three surprise-related conditions we first observed in Fig. 3a. Global population-vector Pearson correlations across trial types are significant for all trial-type pairs (Fig. 3h, top-right; diagonal elements are zeroed and all off-diagonal elements have *p <* 0.01 via permutation test, see Methods for details), and quantify the qualitative contextual differences noted in Fig. 3h, top-left.

By contrast, shuffled controls and stimulus-integrator neurons show relatively little contextual modulation across trial types (Fig. 3i,j, Fig. S4c,d). Thus, contextual modulation is a specific property of the sequence neurons.

Motivated by the resemblance of the properties of these sequence cells to hippocampal representations, we examined the anatomical distribution of sequence neurons relative to stimulus-integrator neurons. We found a disproportionate contribution from entorhinal cortex (ENT), hippocampal subfields, and neocortical regions (Fig. 3k) in the sequence neurons (hippocampal formation 13.7%, isocortex 22.1%) relative to the stimulus and integration populations (hippocampal formation 9.1%, isocortex 15.3%).

Most strikingly, the sequence neurons constitute one of the largest functional categories in the brain during this task: 21.8% of all recorded neurons, compared to 0.8% for stimulus-responsive, 4.5% for integrator, 10.3% for movement-initiation, and 3.7% for movement-execution neurons (Fig. 3l). Although it has recently been appreciated that movement-related signals dominate over sensory-driven responses across the brain ( ^(19;37)^), our data show that internally generated sequential activity is one of the largest identifiable single populations, exceeding all individual classical sensorimotor categories and second only to the combination of all movement-associated categories (including movement-feedback) — at least in expert animals performing a well-learned task (Fig. S3).

In sum, during each trial the brain exhibits a widespread propagation of temporally precise, sparse, and reproducible neural sequences. These sequence neurons are distinguished from all functional categories along many different dimensions. They share a common backbone across trial types but carry contextual information that discriminates hierarchically between surprise and non-surprise conditions and further between individual trial types. Their widespread presence across the brain together with their specific hippocampal-entorhinal bias suggests that they might originate in hippocampus and be spread across the brain and especially to the neocortex to supply abstract scaffolds that support the association of world states with internal states that delineate context and time.

### Low co-correspondence of function with anatomy except cortical-subcortical division

Next, we consider the localization of these functionally specific clusters to anatomical regions in the brain. Thus, we use our brain-wide recordings to address the question of whether basic functions are primarily assigned to specific brain regions that pass their outputs to one or a few downstream areas for the next stage of sequential processing (Fig. 1a, left), or whether the brain follows a broadcast model of computation where each area broadcasts its signal to many different regions, any or all of which might use that signal for parallel computations of their own (Fig. 1a, right).

We assessed localization in several ways: first, we colored each row of the rastermap by that neuron’s Beryl region (Fig. 4a, see methods for anatomical parcellation information). The result is a muted mixture of colors, with the only visible separation given by pink-green color blocks indicating a subcortical (thalamus, hindbrain, midbrain)-cortical (isocortex, hippocampus) divide in neural response patterns. Conversely, we can sort and color neurons by Beryl region, but then lose the functional structure of the spike rasters (Fig. 4b). All the rich functional categories and spatiotemporal response patterns of the brain-wide rastermap are present if we constrain the rastermap to single brain regions with sufficiently many cells (Fig. 4c), whether the regions are in cerebellum, thalamus, isocortex, hippocampus, or midbrain. Thus, the rastermap confirms and demonstrates in detail that individual Beryl areas contain cells showing essentially the full diversity of temporal, trial-phase and trial-type specific response patterns, consistent with the broadcast view of computation.

**Figure 4.**
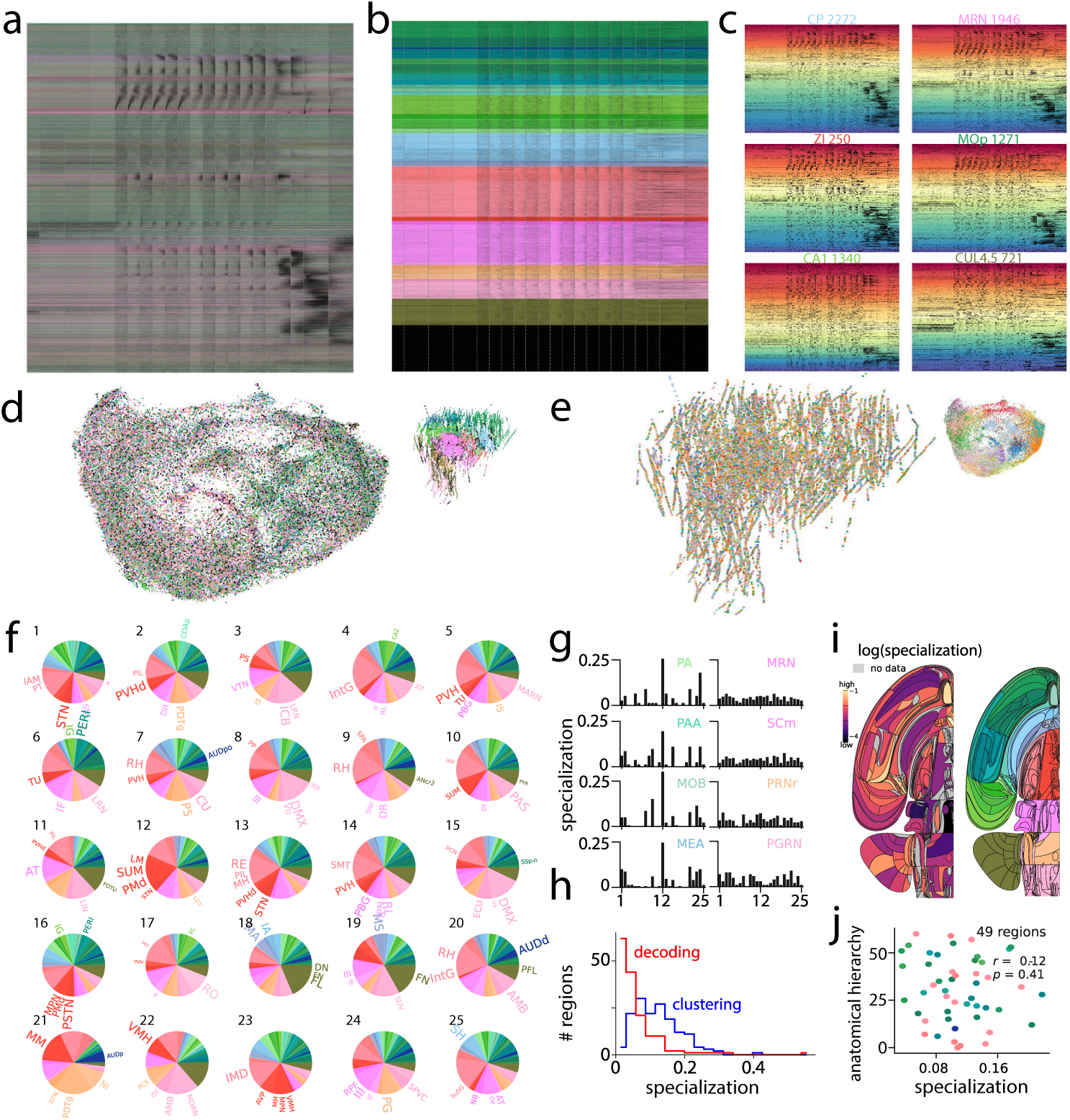
Low functional and anatomical co-correspondence except strong cortical-subcortical division. **a** Response vectors sorted by similarity (rastermap) overlaid with anatomical region colors (Beryl) does not correspond beyond a weak green (cortex, hippocampus) and pink (thalamus, hindbrain, midbrain) divide. **b** Response vectors sorted by anatomical region overlaid with anatomical region colors (Beryl) suggests the presence of most response patterns in most regions, however no more visible functional fine structure. **c** Response vectors for 6 example regions (CP, caudoputamen; MRN, midbrain reticular nucleus; ZI, zona incerta; MOp, primary motor area; CA1, hippocampal field CA1; CUL4,5, cerebellar culmen lobules IV–V), sorted by similarity and overlaid by rastermap cluster colors, showing all main response patterns in each region. **d** 2D embedding (UMAP) of response vectors colored by anatomical region (Beryl) does not show much correspondence beyond a green (cortex) and pink (midbrain/hindbrain) divide, while the 3D positions of the cells in anatomical space match the anatomical region colors by definition (small point cloud). **e** 3D positions of the cells in anatomical space, colored by functional cluster (25 k-means) shows no correspondence, while the 2D embedding (UMAP) of response vectors is visibly divided by functional cluster colors (small point cloud). **f** Regional composition of each functional cluster group of neurons, pie charts sorted canonically, shows that all functional clusters contain most regions with only few outliers such as cluster 21. 5 most numerous regions are labeled in each. **g** Regions differ by specialization. Counts of cells across functional clusters (k-means) for 4 example regions with the least flat distribution (highest specialization) and the 4 most flat ones. See Fig. S5 for all regions. **h** Distribution of specialization scores based on k-means cluster counts across all regions, next to the distribution of specialization scores based on decoding of 4 main IBL variables (results from (^21^)). Specialization via both methods correlate strongly across regions, Fig. S6a. **i** No clear correspondence between specialization scores (left) and anatomical partition (right); Swanson flatmap. **j** Correlation between anatomical hierarchy (based on tract tracing (^39^)) and specialization score based on k-means cluster counts is not significant.

Next, we consider whether the clusters in the UMAP single cell response embedding of Fig. 2g align with Beryl region coloring as they did with k-means prototypes. Instead, we found a near-total lack of alignment between functional clusters and regions (Fig. 4d). Conversely, when we visualize the three-dimensional anatomical position of each recorded cell, colored by its k-means functional cluster, the result is a spatially homogeneous mix (Fig. 4e), in contrast to the previously observed alignment of k-means and UMAP functional clusters (Fig. 2g).

For each functional response category (i.e. k-means cluster average response), we can construct a map of Beryl region membership (Fig. 4f). Here, almost each category is represented in almost all brain regions, with seemingly similar area distributions per category, and some notable exceptions (e.g. categories 21, 22, which visibly depart from the other distributions). We next quantified, for each region, its distribution of functional categories, and found a difference in regional distributions of functional cell types between regions. Different regions have different levels of “specialization” (Fig. 4g): some regions contain a peaky distribution of functional categories, so are specialists (e.g. PA, MOB, MEA), while others are generalists with a relatively equal distribution of cells across functional categories (e.g. MRN, PRNr, SCm; see all regions in Fig. S5). We cross-checked our measure of specialization based on our unsupervised functional categories with an estimate of specialization based on decoding specific variables (results from(^21^)) and found a strong correlation (Fig. S6a) with lower overall specialization through decoding but a similar tail (Fig. 4h). Overall, subcortical regions including midbrain (pink, e.g. MRN) and hindbrain (orange, e.g. PRNr) tend to exhibit lower specialization (Fig. 4i), while those with high specialization tend to be cortical (green, e.g. MOB), earth-mover distance 0.041 between the two distributions (also see Fig.S11 for an alternative statistical quantification of the cortical/sub-cortical divide). We found no other simple relationship between commonly defined brain axes or partitions and specialization, or when we overlay the specialization scores onto the Swanson map (adapted by(^21^) from(^38^)), which showed that specialization does not co-cluster with Cosmos level. To compare these results to the conventional view that lower brain regions are more functionally specialized while higher brain regions are generalists, we directly plot functional specialization scores against a structural hierarchy score obtained for a subset of brain regions via tract-tracing (Fig. 6d in(^39^)). We found that structural hierarchy and specialization are not significantly related, regardless of specialization metric (Fig. 4j and Fig. S6). Thus, we do not find that there is a relationship between regional hierarchy level and the diversity of functional response types contained in the region.

Next, inspired by fMRI studies that investigate functional networks in mouse brain^(40;41;42;43;44;45;10;46;47;48)^, we applied multi-region functional graph analysis methodologies to generate maps of cross-brain correlation during different phases of the task. We defined a similarity score *S_ij_* of neural activity in Beryl regions *i* and *j*, computed as the cosine similarity between 2D embeddings of neural response features (Fig. S9a) in a particular task phase and condition. By applying Louvain community detection to the resulting similarity matrices in selected task phases (i.e. subsets of PETHs, see SI for definition), we identified non-orthogonal functional similarity subnetworks to reconfigure between task phases and conditions (Fig. S9b-e, details in SI).

When we correlated key fMRI-defined functional networks in the mouse brain (such as the default mode network and the sensory mode network; see SI for details) with the functional similarity subnetworks we identified, we found little overlap between fMRI-defined anatomical networks and our functional maps (Fig. S9g). Despite the reconfiguration of functional subnetworks across task phases, one division of subnetwork compositions remained consistent across tasks phases: a cortical-subcortical functional division in the form of a two-block structure in the subnetwork similarity matrix (Fig. S9f). This result reinforces our above finding of a cortical-subcortical division in functional responses (also see S11).

In sum, despite finding clear functional response categories from the level of neural populations down to single neurons, we find low to no regional localization. Cells performing all functions are nearly homogeneously distributed across the brain, with at most a cortical-subcortical distinction, consistent with the existence of distinct but intermingled functional subcircuits within an area, sometimes characterized by genetically defined cell differences or different projection targets or inputs^(49;50;22)^.

Overall, these findings and the changing functional coalitions are consistent with the compute-broadcast-select view of computation in the brain where different neurons in a region participate in different processing loops involving various brain regions, gating and selecting the broadcast inputs they receive, and helping determine the functional allegiance of the region based on relative subnetwork activation in a given task phase.

### Structured, not random, mixed selectivity at the single-neuron level

A prominent hypothesis holds that neurons in association cortex encode random mixtures of task-relevant variables — a property termed random mixed selectivity and motivated by recordings in prefrontal cortex showing that individual neurons responded to conjunctions of multiple task variables rather than single variables in isolation(^15^). The theoretical appeal is that random, high-dimensional representations maximize the number of possible linear readouts, thereby supporting flexible behavior in novel task contexts^(51;52)^. Despite the influence of this idea, a direct test of the independence assumption — against an appropriate null model, and across the brain rather than within a single area — has not previously been performed. The random hypothesis would predict that the features a neuron combines are statistically independent, so that knowing a neuron encodes stimulus identity provides no information about whether it also encodes choice, feedback, or prior expectation.

The combination of interpretable functional categories per neuron (Fig. 2a-c) with brain-wide coverage and large neuron counts places us in a position to perform such a test. We constructed a set of 100 prototype response vectors by k-means clustering of the feature vectors and averaging across cluster (Fig. S7, and see methods), then projected each neuron’s response onto this prototype basis to obtain a 100-dimensional coefficient vector per cell — a compact description of how each neuron combines the identified functional motifs. If mixed selectivity is near-random, these coefficients should be statistically independent across neurons, conditioned on their marginal distributions.

The data clearly violate this prediction: The ordered coefficient matrix, where rows are neurons (ordered by the canonical rastermap ordering) and columns are coefficients (hierarchically ordered based on the Pearson correlation of the 50k-dimensional columns), exhibits prominent block structure, with consecutive groups of neurons sharing highly correlated coefficient profiles (Fig. 5a); we can also see this in the structured nature of the (z-scored) coefficient correlation matrix (Fig. 5b) and the observation that specific features consistently co-occur across neurons in neural groupings (Fig. 5c). Fig. S7 allows to coarsely interpret the co-occurring response categories: coefficients 2-19 strongly co-occur in neurons, mixing primarily choice-related accelerating upward ramps with change-related signals (2,13) and movement-initiation signals (4,11). Coefficients 19-38 strongly co-occur in neurons, mixing primarily movement with quiescence, mistake, and both types of feedback. Neurons mix coefficients 43-47, corresponding to a mixture of block coding with correct feedback. And neurons mix coefficients 50-79, combining narrow activation peaks at different delays (corresponding to fixed-latency localized firing as for sequences) with some linear downward ramping towards movement onset. Finally, coefficients 80-100 form a weaker grouping, and consist of sensory onset coding together with movement decision signals (peaking shortly before movement, but decaying strongly into movement).

**Figure 5.**
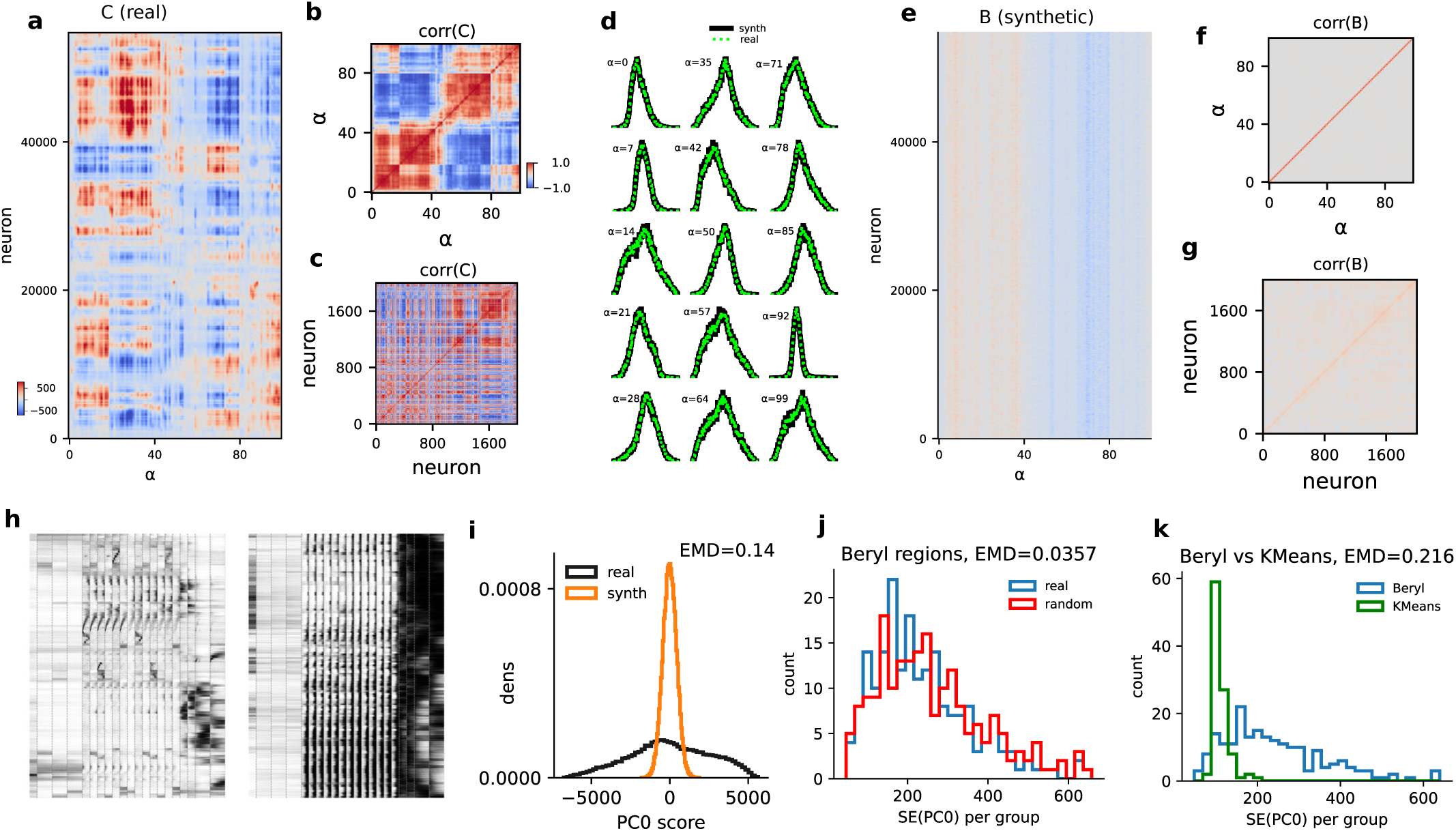
Structured, non-random selectivity at the cellular level with random distributions at the regional level. **a** Projection coefficients *a* of ∼50,000 neurons onto 100 k-means cluster-average response vectors (coefficients sorted hierarchically based on their Pearson correlation matrix; this sorting is used across all relevant panels in this figure; neurons sorted by RM sorting, same as Fig. 2f). The coefficients reveal strong structure and correlations across neurons. **b** Correlation matrix of the 100 coefficients across neurons highlights pronounced dependencies, indicating clustering into discrete response types. **c** Correlation matrix of a randomly sampled subset of 2000 neurons (retaining their ordering from a)). **d** Empirical coefficient distributions (green) closely match those of synthetic neurons (black) constructed to preserve marginal statistics. **e** Synthetic neurons’ coefficient matrix, generated by independently sampling coefficient distributions, exhibit weak correlations. **f** Corresponding correlation matrix confirms the absence of higher-order dependencies by construction. **g** Subset of 2000 synthetic neurons likewise shows minimal structure. **h** Left, Rastermap-sorted response vectors reconstructed from real coefficients reproduce structured activity patterns; right, synthetic reconstructions sorted by Rastermap lack structure. **i** Neural response diversity quantified by the distribution of loadings along the first PC (PC0) of the coefficients of real (black) and synthetic (orange) neurons. **j** Regional response diversity quantified by standard error of coefficients along first PC, within anatomical region groups (Beryl) and random allocations of neurons to groups of the same sizes as the Beryl regions. **k** Distributions of standard error of coefficients along first PC, for neurons within Beryl regions versus neurons within functional groups (100 k-means clusters): functional diversity is much lower within functional than anatomical groups.

To ensure that this structure is not inherited from differences in the marginal distributions for each coefficient, we constructed a null model that preserves the marginal statistics (and thus overall prevalence of each functional feature across the population; Fig. 5d, green) of the coefficients, but destroys the joint structure between coefficients. Specifically, we generated 50,000 synthetic neurons by sampling coefficients independently from the marginals. The resulting synthetic coefficient vectors (Fig. 5e, sorted as panel a) entirely lack the diversity and block structure of the data (Fig. 5a), even though the synthetic population’s sampled marginal coefficients nearly exactly match the data’s coefficient marginals (Fig. 5d, black). Vertical bands in the coefficient matrix reflect differences in the marginal distributions for different coefficients. The coefficient and neuron-neuron correlation matrices (z-scored) reflect the lack of structure in the null model (Fig. 5f-g).

To verify that the prototype basis of the generative model used in our null model is rich enough to reproduce neural responses, we used it to reconstruct real neural responses from their prototype coefficients (Fig. 5h, left). This confirms that the failure of the null model to match the data (Fig. 5e-g, h, right) is due to the departure of data from the independence assumption, rather than an impoverished basis set.

We quantify these results by the distribution of the first principal component of the neural coefficient vector across neurons and synthetic neurons. There is a large difference in these distributions, with a much larger width of the natural distribution compared to the synthetic one (Fig. 5i; earth-mover distance, EMD=0.14), reflecting the much greater diversity of cellular responses in neurons relative to the randomly generated synthetic neurons. Together, these results argue against random mixed selectivity as a general principle of neural coding, at least in the context of a well-learned task.

Finally, though neurons combine features in diverse but structured ways, the spatial distribution of neural features across brain regions is quantifiably much closer to random. The distribution across Beryl regions of the first principal component of the neural coefficient vectors within each region is similar to the distribution when we construct mock Beryl regions by shuffling neurons between them (Fig. 5j, EMD 0.0357; the mock regions have the same neural partition sizes as the Beryl regions). Thus, the distribution across Beryl regions is much more similar to its null distribution than the distribution of single neuron responses is to its null. The Beryl distribution can also be compared to the same distribution constructed for functionally clustered neural groupings rather than Beryl groupings, reflecting the high similarity of functionally grouped neurons (Fig. 5k, EMD 0.216). All the above results are qualitatively unchanged if we vary the number of prototype features being considered from 100 to 40 (Fig. S8).

These results reveal a striking dissociation across representational levels in the brain. At the single-neuron level, selectivity is diverse but structured: neurons combine specific features in highly correlated functional groupings and we find little evidence of random mixed selectivity as a general property of representation in neurons. At the regional level, these structured functional neuron types are distributed across the brain in proportions close to what random allocation would produce. The implication is that the brain’s computational architecture is organized around stable, non-random functional motifs that are homogeneously dispersed across regions, consistent with the anatomical analyses of the preceding section and with the compute-and-broadcast model.

## Discussion

The combination of brain-wide cellular-resolution recordings with a behaviorally rich task structure has allowed us to characterize the functional identity of ∼50,000 individual neurons and examine how these identities are organized across the mouse brain. Three findings, taken together, challenge the prevailing staged model of brain computation and point toward a broadcast architecture structured by internally generated sequential activity.

First, unsupervised clustering reveals that individual neurons fall into distinct and interpretable functional categories, each dominated by a specific aspect of the task: stimulus detection, evidence integration, movement initiation, movement execution, feedback processing, or representation of internal states such as block prior and prediction error. Ordering neurons by the similarity of their responses reveals the temporal arc of these computations at cellular resolution — distinct populations transiently activate to process one aspect of the task, then yield to the next, from stored expectations through perception, decision, and action. Importantly, most individual neurons are dominated by selectivity to a single sub-function within the task rather than exhibiting the broad, heterogeneous tuning often attributed to neurons in association cortex.

This observation bears directly on the mixed selectivity hypothesis. Although neurons do combine features, they do so in highly structured, correlated groupings that depart dramatically from the independence predicted by the random mixed selectivity model^(15;51;52)^. Our null model — which preserves the marginal prevalence of each feature but destroys their joint structure — produces a homogeneous, low-diversity population that is a poor match to the data. Thus, whatever mixing occurs is constrained and specific, not random. One possibility is that the structured co-occurrences we observe reflect the consolidation of task-relevant feature bindings through learning: early in training, representations may be more randomly mixed to support flexible readout, with experience gradually sculpting the correlated groupings we observe in expert animals. This prediction could be tested by applying the same analyses to recordings obtained at earlier stages of training.

Second, despite clear functional specialization at the cellular level, neurons belonging to a given functional category are widely dispersed across brain regions, and each region contains cells spanning nearly the full repertoire of response types. There is thus a bidirectionally low co-correspondence between function and anatomy. The result implies that individual brain regions are not unitary computational modules: each houses multiple local circuits participating in distinct computations, with a given circuit potentially more strongly interconnected with partner sub-circuits in distant regions than with neighboring neurons in the same anatomical region. The computational unit, in this view, is the distributed loop — in the form of cross-regional sub-circuit partnerships — rather than the anatomical region^(53;54;18)^. This view is entirely consistent with the recent understanding of the astonishing diversity of cell types in terms of morphology, connectivity, and gene expression within each region^(23;24;25;26;27;28)^. To this emerging picture, our data add direct functional evidence at single-neuron resolution across the brain that the outputs of these loops are present nearly homogeneously in all regions, as expected if broadcast is continuous and global.

These results extend earlier large-scale electrophysiology studies with substantially greater anatomical sampling. Steinmetz et al. (^55^) sampled neurons across 42 brain regions under the Beryl parcellation during a similar task, concluding that there were separated representations in different regions. By contrast, the present Brain-Wide Map spans 279 Beryl regions. This increase in coverage makes it possible to detect sparsely distributed functional populations that smaller datasets may undersample. For example, some regions previously appeared to contain no choice-related neurons, whereas in the present dataset choice-related responses are detected much more broadly. We therefore interpret these differences not as a contradiction of earlier findings, but as a consequence of increased sampling density and anatomical coverage: more comprehensive brain-wide datasets can reveal qualitatively new organizational structure that is inaccessible at smaller scale.

We note that the observation that functional responses are dispersed does not establish that they are generated in a distributed way. It remains possible, and in our opinion likely, that specific computations may originate in specific local circuits and are then broadcast for downstream use(^13^). Distinguishing broadcast of locally generated signals from distributed computation will require perturbation experiments that selectively silence candidate generator circuits while monitoring brain-wide responses.

The primary regional distinction we found was between cortical and subcortical processing. Integration, choice, and motor initiation are dominated by subcortical neurons, suggesting that for expert performance on a well-learned task, subcortical circuits may carry much of the computational load. This is consistent with the general principle that extensively practiced behaviors progressively become more subcortically controlled^(56;57;58)^ though we cannot rule out a continued cortical role that is masked by redundancy or by the correlational nature of our analyses. Whether the cortical-subcortical balance shifts during learning — with cortex playing a larger role during initial acquisition — is an important question for future work. Simpler processing differences between cortical and subcortical populations, such as blood volume and a type of whisker correlation(^59^) may explain the divide independent of learning.

Despite the overall homogeneity of functional types across regions, some finer regional structure is present. Regions differ in the degree of functional specialization, with some (e.g., PA, MOB, MEA) exhibiting peaked distributions of functional categories and others (e.g., MRN, PRNr, SCm) containing a nearly uniform mix. However, we found no relationship between these distributional specializations and previously defined anatomical hierarchy scores. Examining co-active regions within each task phase revealed distinct patterns of inter-regional correlation that reconfigure rapidly as different computations unfold. These emergent phase-specific functional similarity networks reinforce the cortical-subcortical distinction seen throughout this work, and form a detailed atlas for future studies of brain-wide coordination. Their relatively weak overlap with fMRI-derived brain networks likely reflects fundamental methodological differences. These include large differences in the measurement modality (neural activity versus blood flow), spatiotemporal resolution (millisecond-scale single-cell spiking versus second-scale population responses), and spatial coverage in terms of number of subcortical regions. The divergence warrants further cross-modal investigation, including across tasks, recording techniques, and species.

One caveat of our study is that it is focused on highly trained mice. It is possible that random mixed selectivity exists during initial learning, when flexible readout of novel feature combinations is at a premium, and that the structured co-occurrences we observe reflect the consolidation of task-relevant feature bindings through experience. This possibility could be tested by examining neural responses at earlier stages of training. Additionally, it is possible that our finding of spatially distributed representations is related to the fact that mice have less-specialized brain structures relative to primates^(4;60)^. Large-scale recordings in macaques will help to address this question.

Third, and most striking, one of the largest fractions of brain-wide neural activity during the task consists not of sensory or decision-related signals, but of internally generated temporal sequences. These sequences tile each trial with sparse, precisely timed activation, maintaining near-constant summed population activity — a motif shared with hippocampal time cells^(34;33)^ and HVC premotor neurons in the songbird^(35;61)^. They share a common backbone across trial conditions but carry graded contextual information, discriminating hierarchically between surprise and non-surprise conditions and further between individual trial types. Their disproportionate hippocampal and entorhinal involvement suggests they may originate in the hippocampal complex^(33;62)^ and be broadcast brain-wide, consistent with the architectural principles outlined above. That internally generated sequential activity dominates over sensory-driven signals and is comparable to movement-related signals, which themselves have only recently been recognized as widespread^(19;37)^, reframes the hierarchy of brain-wide neural dynamics: the cognitive or internal states of the brain account for one of the largest shares of neural variance in expert animals performing a well-learned task.

The brain-wide sequences we report invite comparison with sharp-wave ripple (SWR)-associated replay, the best-characterized form of neural sequences in the mammalian brain. Both phenomena involve hippocampal-entorhinal circuits and both propagate activity to distributed brain regions^(63;64;65)^. However, the two differ in several fundamental respects. SWR sequences occur during quiescence, slow-wave sleep, and immobility — offline states in which the hippocampus is thought to consolidate or plan by reactivating compressed representations of past or future trajectories^(66;67;68)^. The sequences we observe unfold during active task engagement, spanning the full interval from trial onset to movement at the timescale of the unfolding cognitive process itself. SWR content typically recapitulates or recombines spatial trajectories from prior experience; our sequences tile trial time de novo on each trial, with a fixed backbone modulated by current context. SWRs are discrete, intermittent events at variable times; our sequences are time-locked to trial onset and present on every trial.

These distinctions suggest a complementary function. While SWR sequences operate offline to consolidate, evaluate, or plan, the trial-locked sequences we observe may provide an online temporal scaffold — an internally generated coordinate system that indexes time and context during ongoing experience. This interpretation is consistent with data showing that entorhinal and hippocampal representations tile traversals through abstract spaces^(34;69;70;71)^, and with proposals that hippocampal circuits generate temporal frameworks to organize experience^(72;73;74)^. Our findings extend these proposals by showing that the resulting scaffold is broadcast brain-wide, potentially providing a common temporal reference frame for distributed computations. Whether online sequences and offline SWR replay draw on overlapping or distinct circuit mechanisms — and whether the same neurons participate in both — remains open for future investigation.

Several limitations should be noted. Our analyses are based on a single task in expert animals; the balance between functional categories, the prevalence of sequential activity, and the degree of regional homogeneity may all differ during learning, in naive animals, or in tasks engaging different cognitive demands. Recordings are restricted to the left hemisphere, so we cannot assess inter-hemispheric organization. Our conclusions are correlational: establishing causal roles for the identified functional categories, and distinguishing local generation from broadcast reception, will require targeted perturbation experiments. Finally, though the supersession approach provides unprecedented brain-wide coverage at cellular resolution, it relies on alignment across animals performing an identical task, and may miss idiosyncratic or animal-specific response patterns.

Together, these findings suggest a revised architecture of brain computation: non-random, interpretable selectivity at the level of single neurons; sub-regional circuits organized into distributed, cross-regional loops that continuously broadcast their outputs brain-wide; and, dominating all other modes of neural activity, internally generated temporal sequences that may scaffold the brain’s representation of time and context during ongoing experience.

## Methods and Materials

### Data and task

We analyzed the brain wide map dataset of the international brain laboratory (IBL), consisting of recordings from 139 mice in 12 labs performing a decision-making task with sensory, motor, and cognitive components, obtained with 699 Neuropixels probe insertions covering 279 brain areas (Fig. S12) in the left forebrain and midbrain and the right hindbrain and cerebellum, including dense sampling across multiple primary visual cortical areas (e.g., VISp and higher-order visual areas; see Tab.5 and(^75^) for the full table). The dataset is publicly available (download instructions; data browsing online tool at https://viz.internationalbrainlab.org) with full details here(^21^). Find also information about histologically obtained anatomical coordinates there.

The International Brain Laboratory (IBL) decision-making task(^29^) is such that on each trial, a visual stimulus appeared on the left or right on a screen, and the mouse had to move it to the center by turning a wheel with its front paws within 60 seconds. The prior probability for the stimulus to appear on the left/right side was constant over a block of trials, at 20/80% (right block) or 80/20% (left block). Blocks lasted for between 20 and 100 trials, drawn from a truncated geometric distribution (mean 51 trials). Block changes were not cued. Stimulus contrast was sampled uniformly from five possible values (100, 25, 12.5, 6.25, and 0%). The 0% contrast trials, when no stimulus was presented, were assigned to a left or right side following the probability distribution of the block, allowing mice to perform above chance by incorporating this prior in their choices. Following a wheel turn, mice received positive feedback in the form of a water reward, or negative feedback in the form of a white noise pulse and a 2 s time-out. The next trial began after a delay, followed by a quiescence period during which the mice had to hold the wheel still. Movement onset is defined as the earliest statistically significant deflection of the steering wheel after the go cue, exceeding an empirical fixed threshold in any direction(^29^).

Anatomical locations were assigned using the Allen Mouse Common Coordinate Framework (CCFv3), a three-dimensional reference atlas and ontology of the adult mouse brain(^76^). We performed most regional analyses using the IBL Beryl mapping, an intermediate-resolution remapping of Allen CCF regions into 308 anatomical groups, introduced for IBL brain-wide electrophysi-ology analyses to reduce fragmentation of the full Allen ontology while preserving regional specificity. Beryl regions are mainly defined at the level of major cortical areas, nuclei, and ganglia, omitting finer laminar and nuclear subdivisions. For coarser anatomical classification, we used the IBL Cosmos mapping, which groups Allen regions into broad anatomical divisions and was used here to define cortical and subcortical classes. For flatmap visualization, we used the Swanson mapping, which relates Allen/IBL regions to the Swanson anatomical hierarchy and flatmap representation^(77;78)^.

### Peri-event time histograms (PETHs)

We binned spiking activity of neurons during trials declared as “good” as in(^21^) with time bins of length 12.5 ms, and stride 2 ms. See data loading functions load_good_units for neural data and load_trials_and_mask for trial data in bwm_loading.py. We decided to choose a set of 21 PETH types (Tab.1) such that a balance is struck between maximizing trial-specific information while keeping as many sessions as possible. I.e. too specific PETH criteria are satisfied by too few sessions. Our choice, with the additional requirement of at least 10 trials for all PETH types, resulted in 569 unique insertions being included in the analysis and grouped together, each neuron being described by a functional feature vector of all 21 PETHs concatenated together, then z-scored to homogenize differences in baseline activity. We used 54719 neurons in the single cell analyses and 160 regions, when restricting to regions with at least 50 neurons, in the regional analyses, and excluding cells that have zero firing rate across all PETHs.

### Umap and rastermap embeddings

We embedded the 50 k cells’ feature vectors (concatenated PETHs, 1872 time bins in total) in 2 dimensions using umap.UMAP (n_neighbors=15, n_components=2)(^79^) implemented in python.

The Rastermap algorithm(^30^) was used in python to sort the 50 k feature vectors by similarity, using these parameters: Rastermap(n_PCs=200, n_clusters=100, locality=0.75, grid_upsample=0, time_lag_window=5, bin_size=1). We visualized the sorted vectors using matplotlib.imshow(data, vmin=0, vmax=1.5, cmap=“gray_r”, aspect=“auto”). Find the full implementation here(^80^).

### Cross-validation on rastermap

To assess the robustness of neural population structure, we implemented an odd/even trial split cross-validation procedure (Fig. S2). Within each trial type and time window, odd-numbered trials were designated as the train set and even-numbered trials as the test set. Trial data within each set were trial-averaged and concatenated following the same procedure described above. The rastermap algorithm ((^30^)) was applied to the train set feature vectors to obtain 100 clusters and a sorting of all 50,000 cells, placing similar cells next to each other. The resulting cell ordering was then used to arrange and visualize the test set feature vectors (Fig. 2f), providing an independent assessment of the spatial and temporal structure identified during training.

### Definition of sequence neurons and other functional types

Functional neuron groups were defined based on their rastermap cluster assignments (Fig. S3). We observe a population of’sequence neurons’ that exhibit consistent but faint sequential activity across all stimulus-onset–aligned PETH trial types in the independent test set raster plot (including change_b_, s, L_s_L_c_L_b_, s, L_s_L_c_R_b_, s, R_s_R_c_R_b_, s, R_s_R_c_L_b_, s, mistake, s), and they roughly correspond to the cells exhibiting prominent diagonal stripe patterns observed in the train set (Fig. S2). Therefore, we used these ‘diagonal stripe’ cells in the train set raster to identify the clusters associated with the faint sequences for analysis (clusters 5-7, 11–20, 45–54, 56–59, in Fig. S3). Stimulus and integrator neurons were identified as cells belonging to cluster 4, and clusters 1-3 and 38, respectively. Movement initiation neurons are in clusters 10, 32, 34-37, 39, 41-42, and 61; movement neurons are in clusters 72-75. Feedback neurons are in clusters 64-66, 78-82, 89, 94-95, and 97. There is an additional group of neurons that activates for both feedback and movement, identified in clusters 0, 62-63, 67-68, 70-71, 76-77, 83-88, 90, and 96.

### Time–time correlation analysis

We performed time–time correlation analyses on sequence and stimulus+integrator neuron groups, exclusively on the test set. For within-condition analyses, we extracted the neurons × time-bins data segment for each stimulus-onset–aligned trial type and computed pairwise Pearson correlations between all time-bin columns (numpy.corrcoef with rowvar=False), yielding a bins × bins matrix that captures the temporal structure of population co-fluctuations. Time bins exhibiting zero variance or non-finite values were masked prior to computation. As a shuffle control, this procedure was repeated after randomly permuting all columns of the PETH matrix independently within each stimulus-onset-aligned trial type and re-applying rastermap sorting algorithm with cross-validation (i.e. applying the algorithm on the train set of trials and using the resulting ordering on the test set of trials). A condition-averaged autocorrelation matrix was additionally computed by averaging PETH segments across conditions before calculating bin-by-bin correlations. All resulting matrices were displayed as heatmaps with the diagonal zeroed.

To characterize the full cross-condition temporal correlation structure, we performed a concatenated bin-by-bin correlation analysis. The PETH matrices across all stimulus-onset–aligned trial types were concatenated along the time axis (neurons × total bins), and Pearson correlation coefficients were computed for every pair of time bins in this concatenated matrix (numpy.corrcoef with rowvar=False). The resulting (total bins) × (total bins) matrix captures both within-condition temporal autocorrelation (on-diagonal blocks) and between-condition temporal cross-correlation (off-diagonal blocks). Time bins with zero variance or non-finite values were masked. A shuffle control was performed by independently permuting the rows within each condition segment independently prior to concatenation, and re-applying rastermap sorting with cross-validation. The resulting matrices were displayed as heatmaps (diagonal zeroed) with condition boundaries indicated by thin black lines and condition labels at block centers. To enhance visibility of weak but structured off-diagonal correlations, a symmetric logarithmic color scale was applied (SymLogNorm, linear threshold = 0.05, base 10), compressing near-zero dynamic range while preserving the sign and relative magnitude of stronger correlations.

### Overall cross-condition correlation

To quantify global similarity between population activity patterns across conditions, we computed pairwise Pearson correlations between all condition pairs by flattening each condition’s PETH segment (neurons × bins) into a single vector and correlating these vectors across all pairs. Statistical significance was assessed via permutation test (1,000 iterations): for each condition pair, the neuron rows of one condition’s segment were randomly permuted to generate a null distribution of correlation values, and two-sided p-values were computed as the fraction of null correlations with absolute value at least as large as the observed value, *p* = (1 + count)∕(1, 000 + 1). The resulting cross-condition correlation matrix is displayed as a heatmap with the diagonal zeroed; all off-diagonal entries reach statistical significance at p < 0.01. This analysis was performed on the test set only.

### Two-dimensional cross-correlogram across trial types

To jointly characterize the spatial (neuron-axis) and temporal (time-bin-axis) similarity of population activity patterns across conditions, we computed two-dimensional cross-correlograms between all pairs of stimulus-onset–aligned PETH segments. Each PETH matrix (neurons × time bins) was treated as a two-dimensional image, mean-subtracted, and smoothed with a 2D Gaussian filter (sigma = 1 along the neuron axis, sigma = 2 along the time-bin axis). Cross-correlations between all condition pairs were computed via FFT-based convolution (scipy.signal.fftconvolve, mode=’full’), yielding full cross-correlograms spanning all neuron and time-bin lags. Each cross-correlogram was normalized by its zero-lag (center) value, such that autocorrelations peak at 1.0 and cross-condition values are expressed relative to this self-similarity. The resulting trial-type × trial-type grid of 2D cross-correlograms was displayed as a matrix of heatmaps with a symmetric logarithmic color scale to improve visibility of weaker off-diagonal structure. This analysis was performed on the test set only.

### Mixed selectivity analysis

Each of the *N* = 54,719 neurons is described by a feature vector **x***_i_* ∈ ℝ*^T^* (*T* = 1,872 time bins) obtained by concatenating trial-averaged PETHs across all 21 conditions and z-scoring. Stacking these vectors yields the feature matrix **X** ∈ ℝ*^N^*^×^*^T^*. Before projection, rows of **X** are permuted by the rastermap sort index ***σ***, i.e. **X** ← **X**[***a***, ∶], where ***σ*** is obtained by running the Rastermap algorithm(^30^) on the full feature matrix with parameters n_PCs=200, n_clusters=100, locality=0.75. The resulting row order places neurons with similar temporal response profiles in adjacent rows.

We applied *k*-means clustering (*k* = *M* = 100) to **X**, obtaining cluster assignments *t* ∈ {0,…*, M* − 1}. Each prototype vector is the cluster centroid

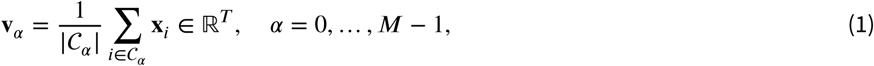

where *C_a_* = {*i* ∶ *t* = *a*}. The prototypes are collected row-wise into the basis matrix **V** ∈ ℝ*^M^*^×^*^T^*. Each neuron’s feature vector is projected onto the prototype basis via an inner product:

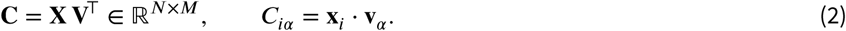

Because **V** is not orthonormal, this is an oblique (non-orthogonal) projection: the entry *C_iα_* measures the raw inner product between neuron *i*’s response and prototype *α*, not the coordinate in an orthonormal basis.

Columns of **C** (basis vectors *α*) are reordered for display by hierarchical clustering of their pairwise Pearson

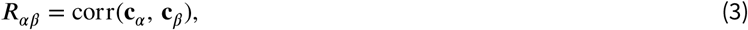

where **c***_a_* ∈ ℝ*^N^* is the *α*-th column of **C**. The distance matrix *D_af)_* = max(0, 1 − *R_af)_*) is passed to average-linkage agglomerative clustering; the leaf order ***,r*** = leaves_list(*Z*) defines the column permutation applied to **C** for all visualisations.

To test the random mixed selectivity hypothesis, we constructed a synthetic coefficient matrix **B** ∈ ℝ*^N^*^×^*^M^* that preserves the marginal distribution of each coefficient independently while destroying all inter-coefficient dependencies. For each *α*, the empirical distribution of {*C_ia_* }*^N^* was estimated with a histogram of 200 equal-width bins over [min*_i_C_ia_*, max*_i_ C_ia_*], giving normalised probabilities {*p_a,b_*}. Synthetic coefficients were then drawn independently:

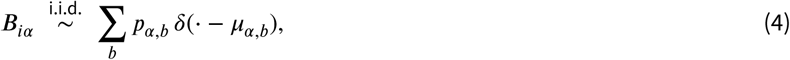

where *µ_α,b_* denotes the centre of bin *b*. By construction, the columns of **B** match the marginal histograms of **C** (Fig. 5d) while their joint distribution is fully independent across *α*. Synthetic feature vectors are reconstructed as

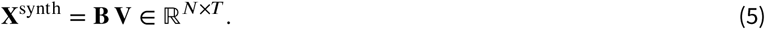

The real and synthetic populations were compared via (i) visual inspection of **C** and **B** and their neuron×neuron Pearson correlation matrices (evaluated on 2,000 randomly sampled neurons); (ii) the distribution of scores along the first principal component of **C** (PC0), quantified by earth-mover distance (EMD = 0.14 between real and synthetic); and (iii) the standard error of PC0 scores within anatomical groups (Beryl regions, size-matched random mock regions, and *k*-means functional clusters), with EMD used to compare group-variance distributions. All analyses used cv=False; results were qualitatively unchanged for *M* = 40.

## Consortia

### The International Brain Laboratory

Larry Abbott, Luigi Acerbi, Valeria Aguillon-Rodriguez, Mandana Ahmadi, Jaweria Amjad, Dora Angelaki, Jaime Arlandis, Zoe C. Ashwood, Kush Banga, Hailey Barrell, Hannah M. Bayer, Brandon Benson, Julius Benson, Jai Bhagat, Dan Birman, Niccolò Bonacchi, Kcenia Bougrova, Julien Boussard, Sebastian A. Bruijns, E. Kelly Buchanan, Robert Campbell, Matteo Carandini, Joana A. Catarino, Fanny Cazettes, Gaelle A. Chapuis, Anne K. Churchland, Davide Crombie, Yang Dan, Felicia Davatolhagh, Peter Dayan, Sophie Denève, Eric E. J. DeWitt, Tatiana Engel, Michele Fabbri, Mayo Faulkner, Robert Fetcho, Ila Fiete, Charles Findling, Laura Freitas-Silva, Surya Ganguli, Berk Gercek, Naureen Ghani, Ivan Gordeliy, Laura M. Haetzel, Kenneth D. Harris, Michael Hausser, Naoki Hiratani, Sonja Hofer, Fei Hu, Felix Hubert, Julia M. Huntenburg, Cole Hurwitz, Anup Khanal, Christopher S. Krasniak, Sanjukta Krishnagopal, Michael Krumin, Debottam Kundu, Agnès Landemard, Christopher Langdon, Christopher Langfield, Inês C. Laranjeira, Peter Latham, Petrina Lau, Hyun Dong Lee, Ari Liu, Zachary F. Mainen, Amalia Makri-Cottington, Hernando Martinez-Vergara, Brenna McMannon, Isaiah McRoberts, Guido T. Meijer, Maxwell Melin, Leenoy Meshulam, Kim Miller, Nathaniel J. Miska, Catalin Mitelut, Zeinab Mohammadi, Thomas Mrsic-Flogel, Masayoshi Murakami, Jean-Paul Noel, Kai Nylund, Farideh Oloomi, Alejandro Pan Vazquez, Liam Paninski, Sabrina Perrenoud, Alberto Pezzotta, Samuel Picard, Jonathan W. Pillow, Alexandre Pouget, Carolina Quadrado, Pranav Rai, Georg Raiser, Florian Rau, Cyrille Rossant, Noam Roth, Nicholas A. Roy, Kamron Saniee, Rylan Schaeffer, Michael M. Schartner, Yanliang Shi, Karolina Z. Socha, Cristian Soitu, Nicholas A. Steinmetz, Karel Svoboda, Marsa Taheri, Charline Tessereau, Matthew Tucker, Anne E. Urai, Erdem Varol, Shuqi Wang, Miles J. Wells, Steven J. West, Matthew R. Whiteway, Charles Windolf, Olivier Winter, Ilana Witten, Lauren E. Wool, Zekai Xu, Kenneth Yang, Yaxuan Yang, Han Yu, Anthony M. Zador and Yizi Zhang.

### Consortium member affiliations

- Center for Computational Neuroscience, University of Washington, Seattle, WA, USA

Leenoy Meshulam

- Center for Neural Science, New York University, New York, NY, USA

Dora Angelaki, Julius Benson, Isaiah McRoberts, Jean-Paul Noel

- Champalimaud Center for the Unknown, Lisboa, Portugal

Jaime Arlandis, Niccolò Bonacchi, Kcenia Bougrova, Joana A Catarino, Fanny Cazettes, Davide Crombie, Eric EJ DeWitt, Laura Freitas-Silva, Inês C Laranjeira, Zachary F Mainen, Guido T Meijer, Pranav Rai, Georg Raiser, Florian Rau, Michael M Schartner, Olivier Winter

- Cognitive Psychology Unit, Institute of Psychology and Leiden Institute for Brain and Cognition, Leiden University, Leiden, Netherlands

Anne E Urai

- Cold Spring Harbor Laboratory, Cold Spring Harbor, NY, USA

Valeria Aguillon-Rodriguez, Cristian Soitu, Anthony M Zador

- Cold Spring Harbor Laboratory, Cold Spring Harbor, NY, USA, Watson School of Biological Science, Cold Spring Harbor, NY, USA

Christopher S Krasniak

- Department of Molecular and Cell Biology, University of California, Berkeley, CA, USA

Yang Dan, Fei Hu

- Department of Applied Physics, Stanford University, Stanford, CA, USA

Brandon Benson, Surya Ganguli

- Department of Basic Neuroscience, University of Geneva, Geneva, Switzerland

Luigi Acerbi, Gaelle A Chapuis, Charles Findling, Berk Gercek, Felix Huber, Alexandre Pouget

- Department of Biological Structure, University of Washington, Seattle, WA, USA

Hailey Barrell, Dan Birman, Kim Miller, Kai Nylund, Noam Roth, Nicholas A Steinmetz, Matthew Tucker, Kenneth Yang

- Department of Brain and Cognitive Sciences, Massachusetts Institute of Technology, Cambridge, MA, USA

Ila Fiete, Ari Liu, Rylan Schaeffer

- Department of Neurobiology, University of California, Los Angeles, CA, USA

Anne K Churchland, Felicia Davatolhagh, Anup Khanal, Maxwell Melin

- Department of Physiology, University of Yamanashi, Kofu, Yamanashi, Japan

Masayoshi Murakami

- Département D’études Cognitives, École Normale Supérieure, Paris, France

Sophie Denève, Ivan Gordeliy

- Gatsby Computational Neuroscience Unit, University College London, London, UK

Mandana Ahmadi, Jaweria Amjad, Naoki Hiratani, Sanjukta Krishnagopal, Peter Latham, Alberto Pezzotta, Zekai Xu

- Institute of Neurology, University College London, London, UK

Kush Banga, Jai Bhagat, Mayo Faulkner, Kenneth D Harris, Michael Krumin, Samuel Picard, Carolina Quadrado, Cyrille Rossant, Miles J Wells, Lauren E Wool

- Institute of Opthalmology, University College London, London, United Kingdom

Matteo Carandini, Agnès Landemard, Karolina Z Socha

- Max Planck Institute for Biological Cybernetics, Tübingen, Germany

Sebastian A Bruijns, Peter Dayan, Julia M Huntenburg, Debottam Kundu, Farideh Oloomi, Charline Tessereau

- Princeton Neuroscience Institute, Princeton University, Princeton, NJ, USA

Zoe C Ashwood, Tatiana Engel, Robert Fetcho, Laura M Haetzel, Christopher Langdon, Brenna McMannon, Zeinab Moham-madi, Alejandro Pan Vazquez, Jonathan W Pillow, Nicholas A Roy, Yanliang Shi, Ilana Witten

- Sainsbury-Wellcome Centre, University College London, London, UK

Robert Campbell, Naureen Ghani, Sonja Hofer, Hernando Martinez-Vergara, Nathaniel J Miska, Thomas Mrsic-Flogel, Steven J West, Yaxuan Yang

- The Allen Institute for Neural Dynamics, Seattle, Washington, USA

Karel Svoboda

- University of California, Los Angeles

Marsa Taheri

- Wolfson Institute of Biomedical Research, University College London, London, United Kingdom

Michael Hausser, Petrina Lau, Amalia Makri-Cottington, Sabrina Perrenoud

- Zuckerman Institute, Columbia University, New York, NY, USA

Larry Abbott, Hannah M Bayer, Julien Boussard, E Kelly Buchanan, Michele Fabbri, Cole Hurwitz, Christopher Langfield, Hyun Dong Lee, Catalin Mitelut, Liam Paninski, Kamron Saniee, Erdem Varol, Shuqi Wang, Matthew R Whiteway, Charles Windolf, Han Yu, Yizi Zhang

## Acknowledgments

This work was supported by grants from the Wellcome Trust (216324), the Simons Foundation, The National Institutes of Health (NIH U19NS12371601) and the National Science Foundation (1707398). We are most grateful for important discussions with Nick Steinmetz.

## Supplementary Material

**Figure S1.**
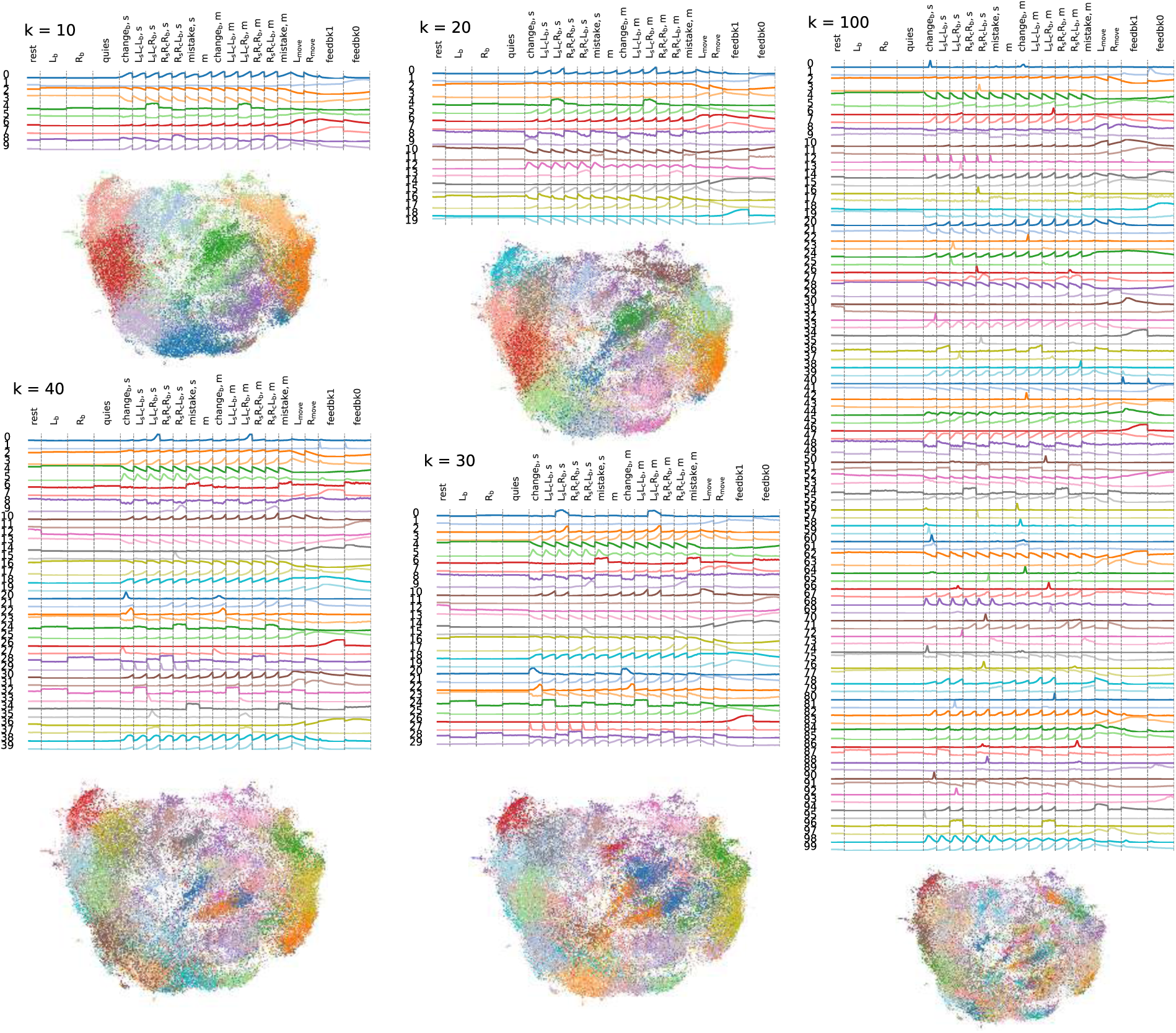
UMAP organization and cluster-mean response profiles across k-means resolutions. For each clustering resolution (*n*_clus_ = 10, 20, 30, 40, 100), the top panel shows the mean trial-aligned response profile of each k-means cluster, and the bottom panel shows the corresponding 2D UMAP embedding of all neurons, colored by cluster identity. The cross-validated data was used here, i.e. the test half of the trials. This figure illustrates how embedding structure and average temporal response motifs evolve with increasing clustering granularity.

**Figure S2.**
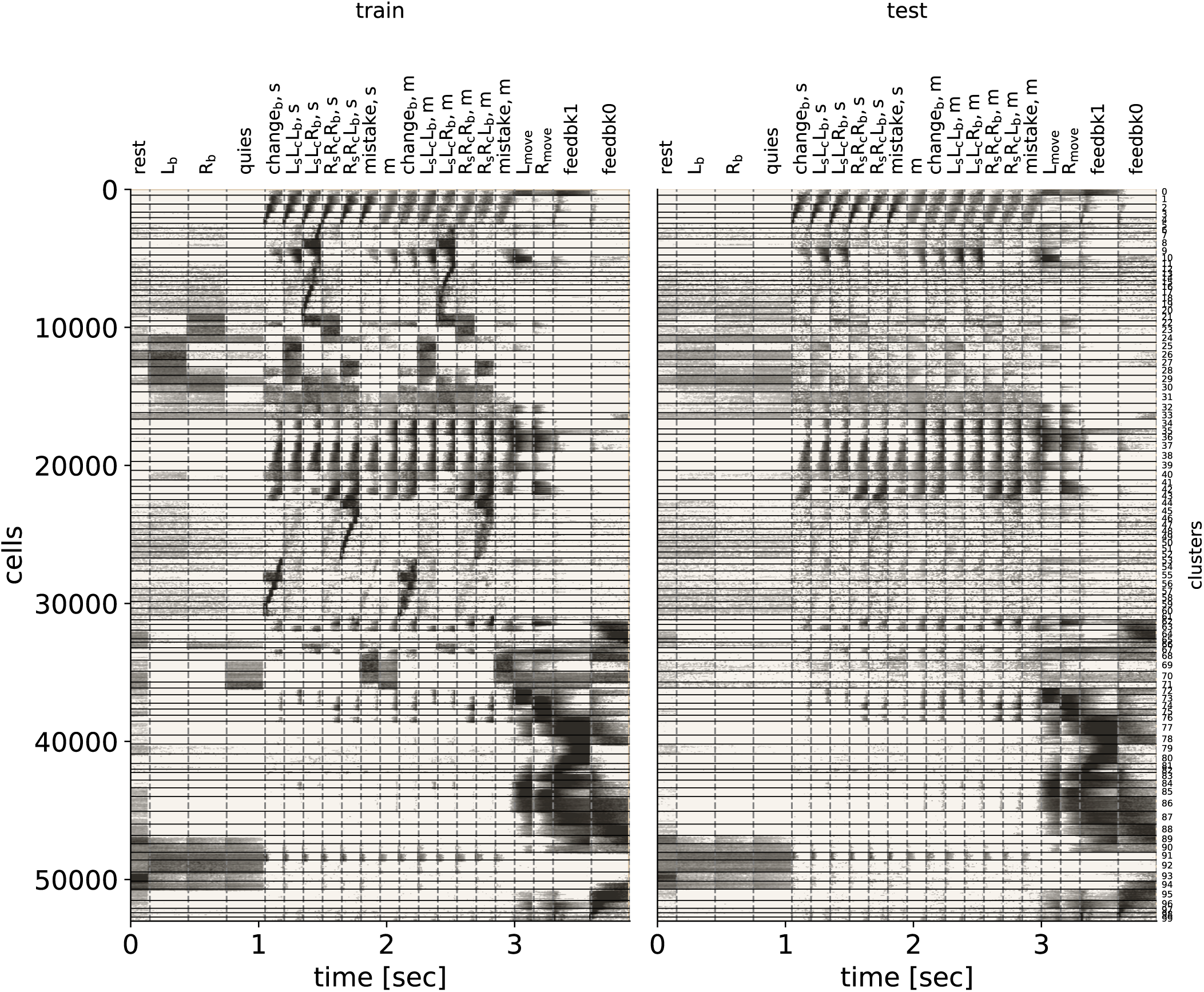
Split-trial cross-validation of rastermap. Trials are split into two sets (train and test) for each alignment and trial type, from which two concatenated feature vectors are computed for each cell. We apply rastermap algorithm on the training set, and apply the resulting ordering and cluster labels to both the train and test set. The majority of the patterns are preserved after cross-validation.

**Figure S3.**
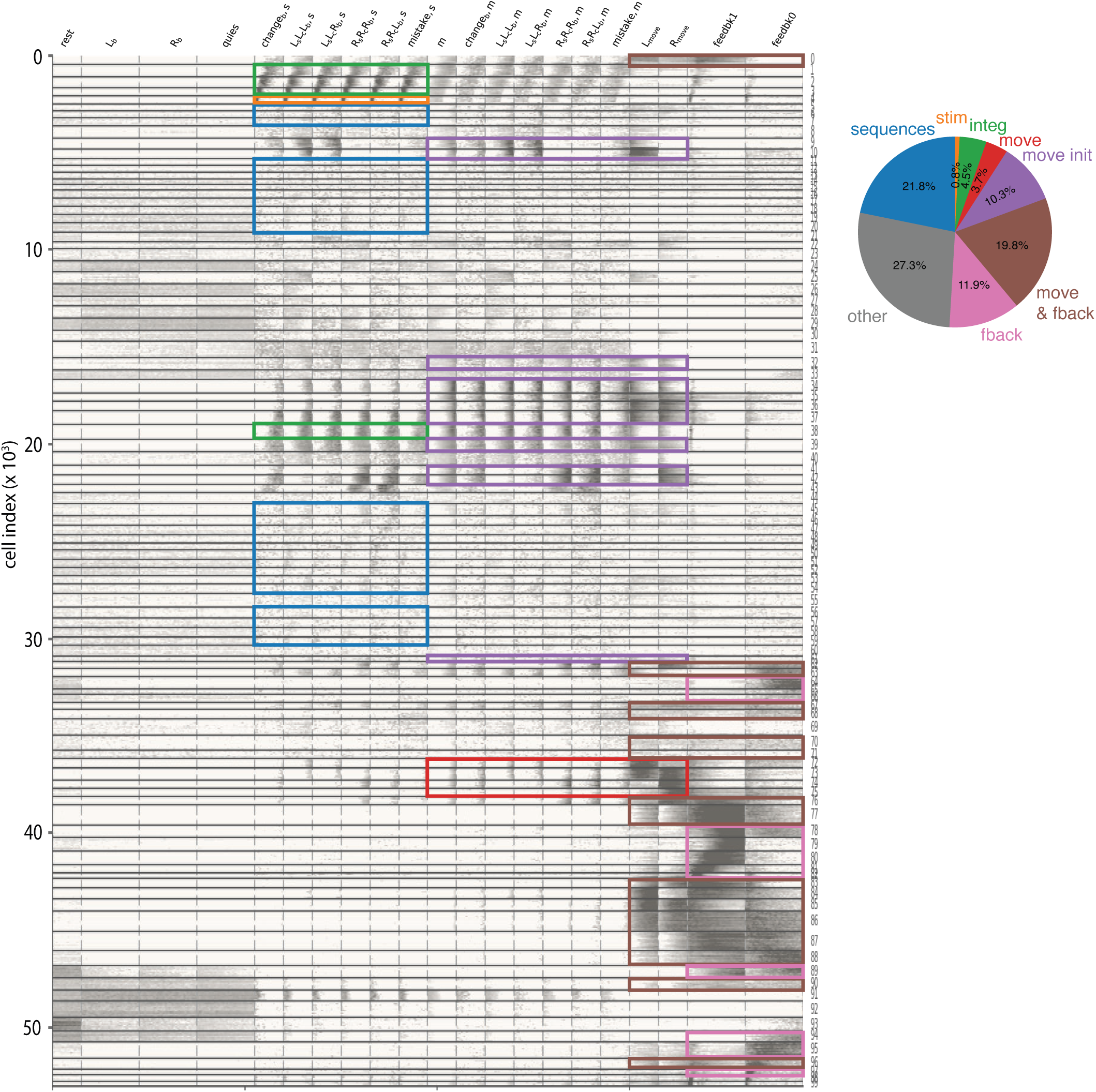
Functional cell clusters identified by rastermap and identification of some simple response categories. Left: Identification of some rastermap categories based on their response patterns: Stimulus response (orange; cluster 4), integrator (green; cluster 1-3, and 38), movement initiation (purple; cluster 10, 32, 34-37, 39, 41-42, and 61) and execution (red; cluster 72-75), sequences (blue; cluster 5-7, 11–20, 45–54, 56–59), feedback (pink; cluster 64-66, 78-82, 89, 94-95, and 97), and co-activation of feedback and movement (brown; cluster 0, 62-63, 67-68, 70-71, 76-77, 83-88, 90, and 96). Stimulus responders are classified as those with sharp early peaks when stimulus-algined, including during the visual stimulus onset and feedback period onset (e.g. for auditory responses). Stimulus integrator neurons are identified as active following the stimulus, with narrower activity packets when stimulus-aligned than movement aligned, with activity decay shortly before the movement period. Movement-related categories are identified based on a strong peak upon movement alignment relative to stimulus alignment. Different movement-related categories are based on the timing of response onset prior to movement, and on whether there is a continuation of activity into the movement period. Right: The corresponding neuron fraction of each group.

**Figure S4.**
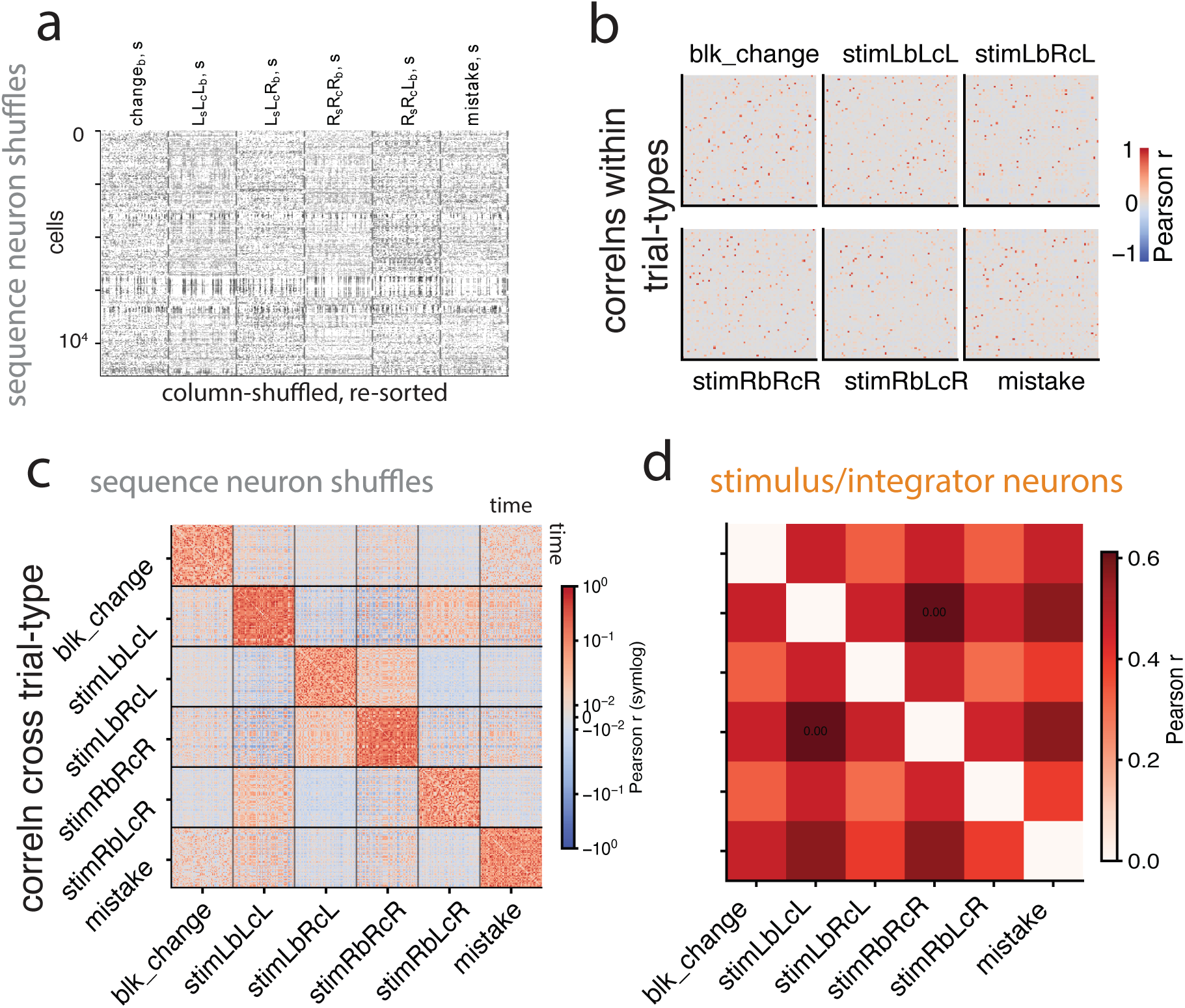
Additional shuffle control for sequence cells. **a** We randomly shuffle the columns (time indices) of the sequence cells independently within each trial type, and re-sort the matrix with rastermap algorithm on the train set of trials (not shown) to apply the resulting ordering on the test set. **b** Time-time correlation matrices of column-shuffled and re-sorted sequence neuron PETHs within each of the trial types, for the cross-validated test set of trials. **c** Time-time autocorrelation across all stimulus-aligned trial types of the column-shuffled and re-sorted sequence cell activity. **d** Overall pairwise Pearson correlation across all stimulus-aligned trial types for the stimulus/integrator neurons.

**Figure S5.**
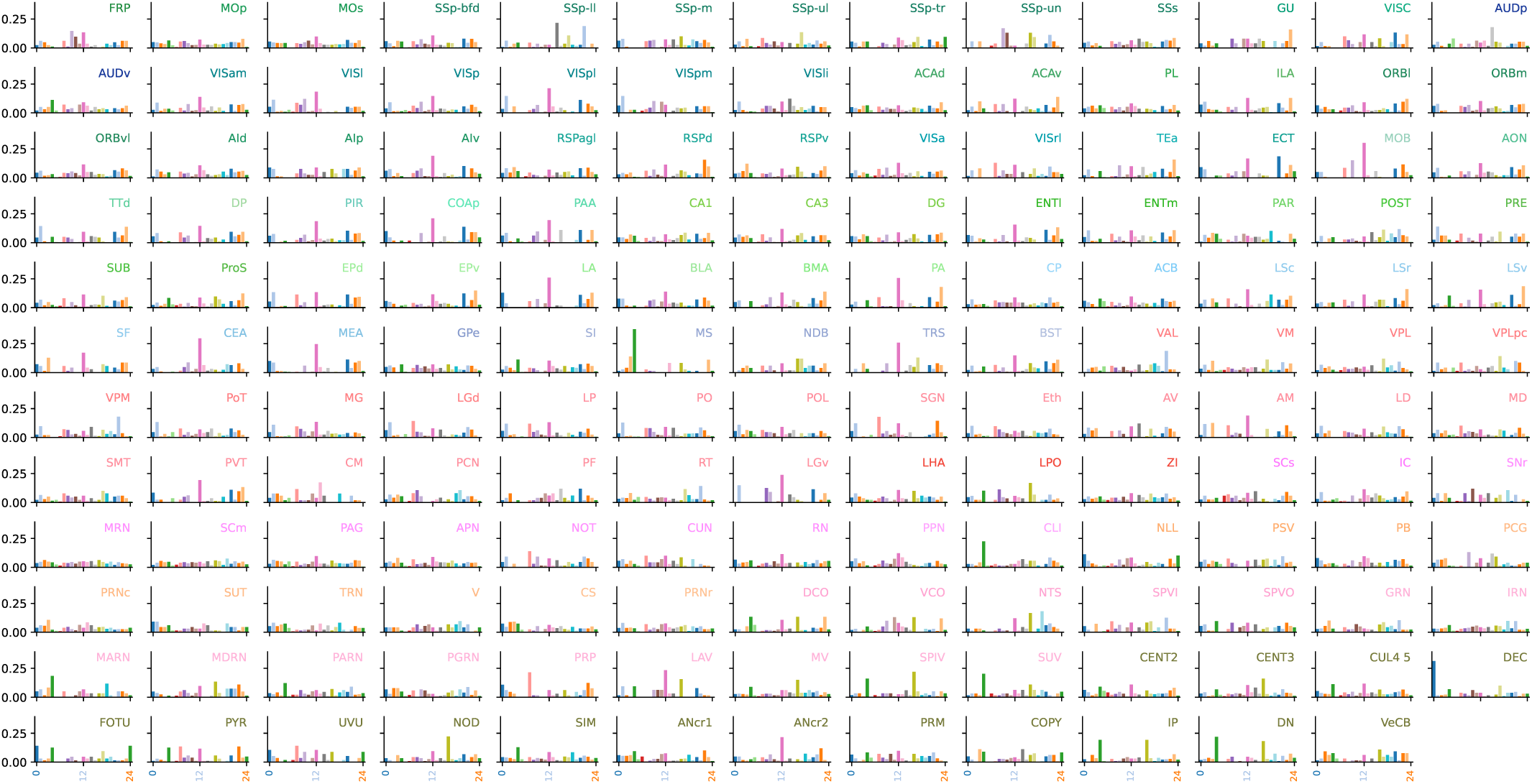
Cluster composition per brain region. Each panel shows the normalized fraction of neurons belonging to each of the 25 k-means clusters for a given Beryl brain region. Only regions with at least 50 neurons are shown (155 regions total). Colors correspond to k-means cluster identity. X-axis: cluster index (0–24); Y-axis: fraction of neurons in that cluster. All regions contain neurons from all clusters.

**Figure S6.**
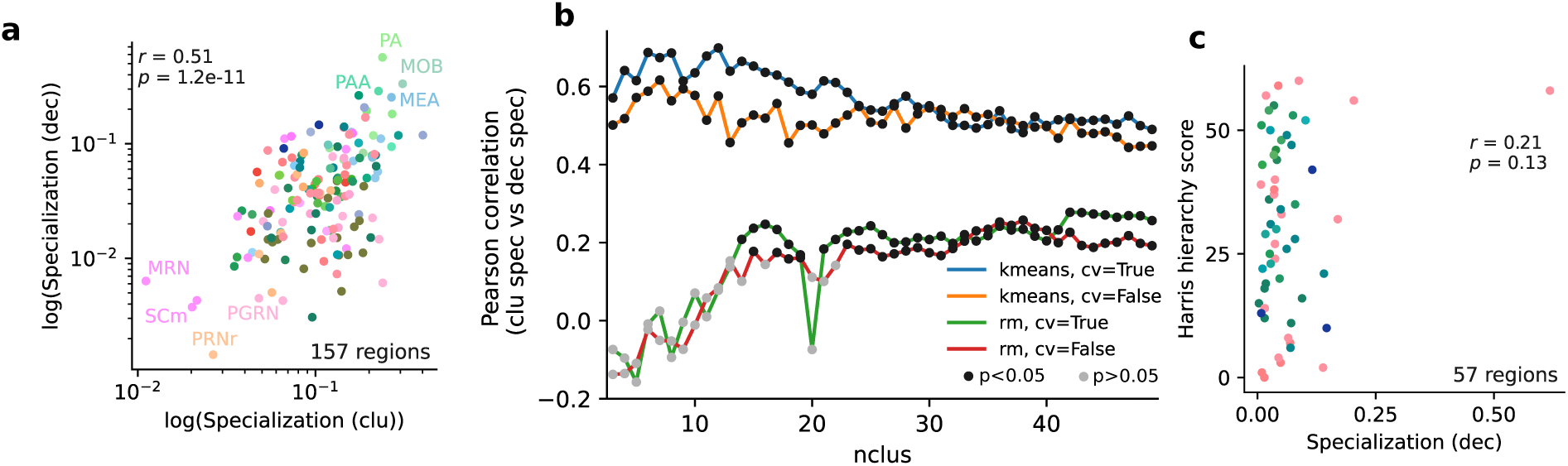
Specialization metrics compared. **a** The k-means specialization score (with 25 k-means clusters) correlates strongly with a decoding-based specialization score. **b** The correlation between the two measures is stable under k-means cluster number variation after a minimal number of about 15 and is also significant when using the clusters provided by rastermap (green, red). Rastermap and k-means results converge as the number of clusters increases. **c** As with the k-means based specialization score in the main text, there is no correlation between decoding-based specialization and anatomical hierarchy score.

**Figure S7.**
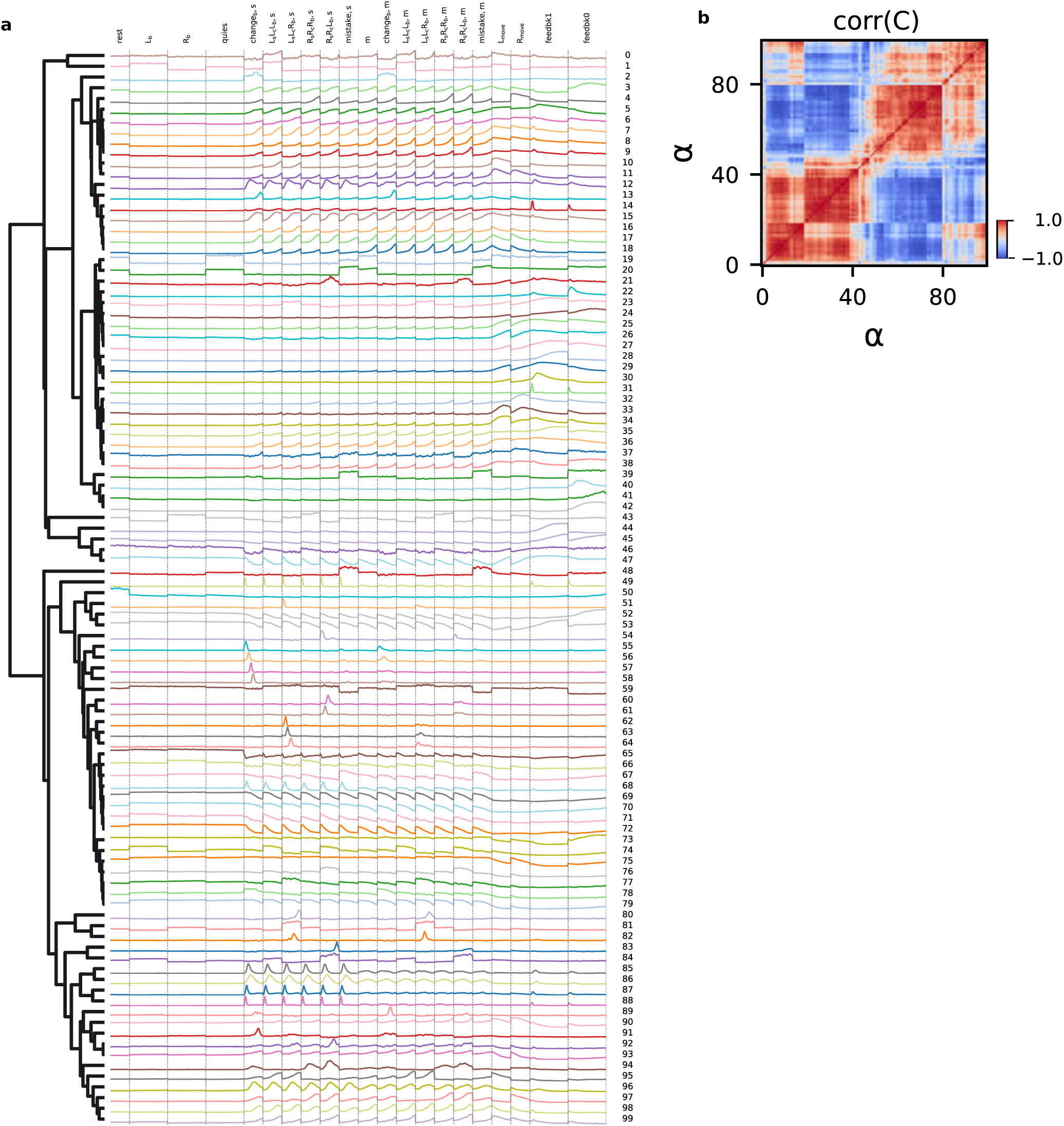
Average feature vectors for 100 k-means clusters, sorted hierarchically. **a** The vectors are shown in order as used in Fig 5a when hierarchically sorting the coefficients of these 100 “basis” vectors (using Pearson correlation as the similarity metric), used to reconstruct the full response data. **b** In the same order, Pearson correlation matrix of coefficients across neurons, as in Fig 5b.

**Figure S8.**
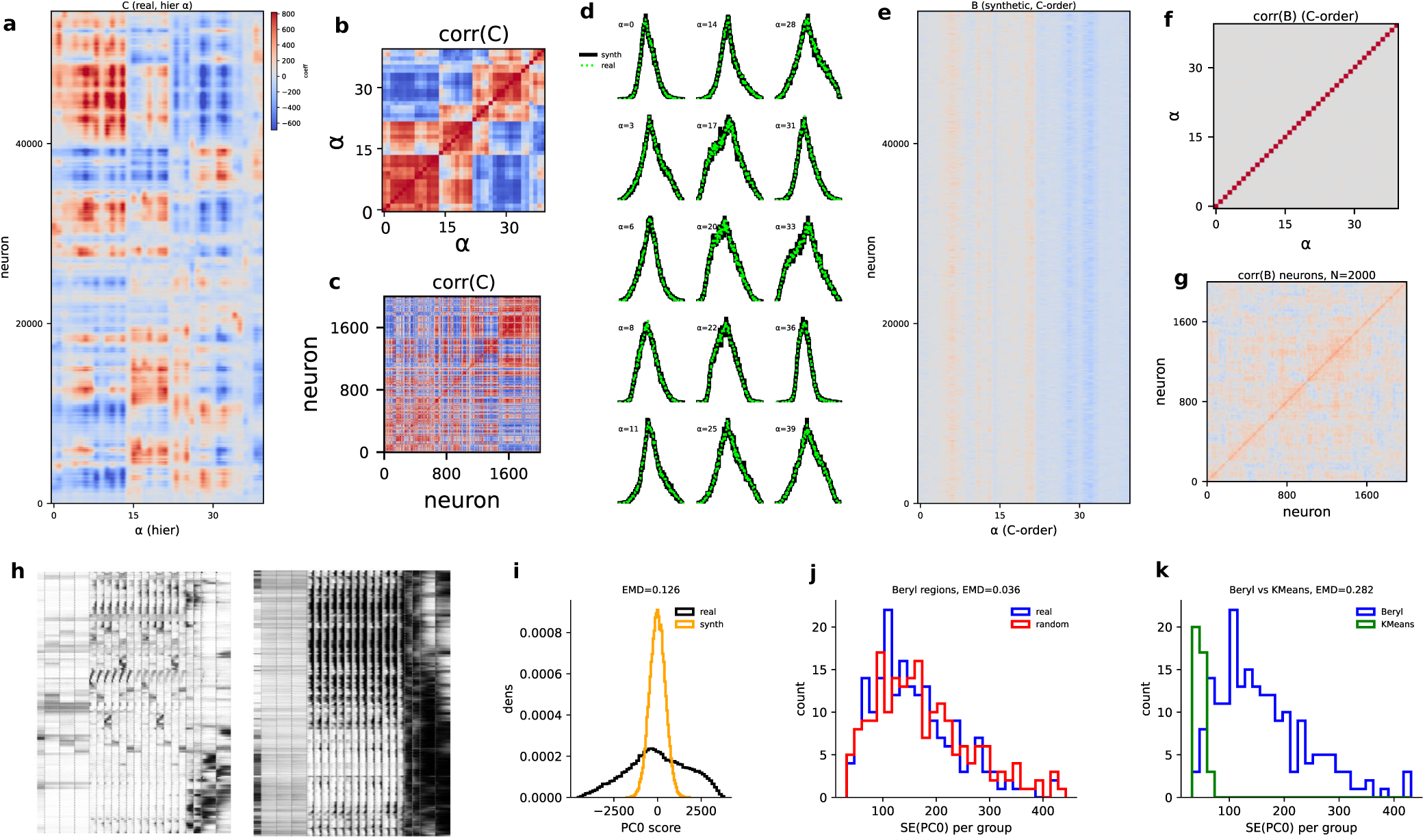
Mixed selectivity analysis for 40 k-means clusters. Same figure as Fig 5 but with 40 k-means clusters as a basis (instead of 100 used in the main figure).

### Dynamically evolving phase-specific functional similarity networks

We have seen that distinct computations unfold in different task phases and involve partially separate neuron groups. A natural question is how different parts of the brain coordinate during these distinct phases. The field of fMRI, with a longer history of brain-wide recordings, has approached this question by performing multi-region functional graph analyses to identify functional subnetworks. The IBL dataset enables similar analyses, but at much higher (∼ 10^3^× or ∼ 1 ms versus ∼ 1 s) temporal and spatial (∼ 10^3^×, with ∼ 60*K* cellular measurements in the supersession versus ∼ 60 voxels) resolution. We now seek an area-wise analysis of dynamical functional similarity networks, or how the brain forms subnetworks to compute over different phases of the task, at high precision.

To assay functional network-level interactions, we defined a functional similarity score between (Beryl) regions per task phase as a proxy for the strength between them, focusing on the following conditions: the resting period (“rest”); the enforced quiescent period (“quies”) and pre-stimulus prior representation in this period (“L_b_” and “R_b_”); the stimulus response period in congruent trials (where the stimulus side equals the block side, i.e. “L_s_L_c_L_b_, s”, “R_s_R_c_R_b_, s”); the stimulus response period in surprise trials (when the stimulus side is opposite the block side, i.e. “L_s_L_c_R_b_, s”, “R_s_R_c_L_b_, s”); the stimulus response period in mistake trials (“mistake, s”); during movement initialization (’m’); in the movement period (“L_move_” and “R_move_”); and during feedback after correct (“feedbk1”) and wrong (“feedbk0”) trials. For each task phase, we subset the z-scored cellular feature vectors (as in Fig. 1f) to the relevant PETH time bins and trial conditions, construct region-wise smoothed 2D activity maps from those phase-specific cellular responses, and compute cosine similarity between the resulting region maps (see the following Method section and Fig. S9a), which yielded a similarity matrix across regions per task phase. All analyses use z-scoring normalization applied before subsetting to reduce baseline firing-rate differences and focus the comparison on response-pattern similarity. Thus similarity matrices compare regions within the same task phase using a common framework, ensuring comparability across phases. We then applied Louvain community detection to the similarity matrices to identify functional networks (region clusters with similar responses), and sorted neurons to be adjacent if they were assigned to the same community, Fig. S9b-h (bottom-left in each panel). Note that the derived networks reflect phase-specific similarity of regional response motifs, not functional connectivity or direct inter-regional coupling. To assess which of the identified networks were reliable and meaningful, we applied three types of control analyses (see the following Methods section), as follows. Only networks that are deemed significant by the intersection of the three controls below are numbered in Fig. S9b-h.

First, we identified which networks had a high’network stability’ score through split-trial comparisons (Fig. S10). We repeated Louvain clustering across several trial splits; for each identified cluster (community) in each split, we defined a binary regional participation vector (whether the region was in the community), and used these to create a regional membership similarity matrix across all clusters and all splits for a condition. We assigned a network stability score (∈ [0, 1]) defined by the average value of its regional membership similarity score (Table 3), and only communities with a stability score exceeding a threshold were deemed statistically stable.

**Table 3.** Stability scores for regional clusters. . Stability scores identified from comparing clusters that are identified from different trial splits for all trial phases. Size indicates the number of times a community is identified in the 8 leave-half-out trial splits analysis. Clusters with stability scores *>* 0.57 and size ≥ 6 (marked as red) pass the stability criteria.

| Cluster | Size | Stability |
| --- | --- | --- |
| concat cluster 0 | 8 | 0.866 |
| concat cluster 1 | 8 | 0.705 |
| concat cluster 2 | 7 | 0.678 |
| concat cluster 3 | 11 | 0.107 |
| resting cluster 0 | 2 | 0.453 |
| resting cluster 1 | 2 | 0.39 |
| resting cluster 2 | 2 | 0.383 |
| resting cluster 3 | 2 | 0.333 |
| quiescence cluster 0 | 7 | 0.65 |
| pre-stim prior cluster 0 | 8 | 0.91 |
| pre-stim prior cluster 1 | 8 | 0.886 |
| mistake cluster 0 | 8 | 0.785 |
| mistake cluster 1 | 8 | 0.624 |
| mistake cluster 2 | 8 | 0.69 |
| mistake cluster 3 | 8 | 0.134 |
| stim surprise cluster 0 | 6 | 0.721 |
| stim surprise cluster 1 | 4 | 0.638 |
| stim surprise cluster 2 | 3 | 0.7 |
| stim surprise cluster 3 | 2 | 0.592 |
| stim congruent cluster 0 | 6 | 0.788 |
| stim congruent cluster 1 | 9 | 0.482 |
| stim congruent cluster 2 | 8 | 0.528 |
| stim congruent cluster 3 | 18 | 0.088 |
| motor init cluster 0 | 8 | 0.899 |
| motor init cluster 1 | 8 | 0.836 |
| motor init cluster 2 | 8 | 0.759 |
| motor init cluster 3 | 12 | 0.153 |
| movement cluster 0 | 7 | 0.839 |
| movement cluster 1 | 6 | 0.729 |
| movement cluster 2 | 8 | 0.661 |
| movement cluster 3 | 15 | 0.083 |
| fback correct cluster 0 | 8 | 0.564 |
| fback correct cluster 1 | 8 | 0.516 |
| fback correct cluster 2 | 7 | 0.523 |
| fback correct cluster 3 | 7 | 0.509 |
| fback correct cluster 4 | 10 | 0.057 |
| fback incorrect cluster 0 | 8 | 0.617 |
| fback incorrect cluster 1 | 8 | 0.602 |
| fback incorrect cluster 2 | 22 | 0.03 |
| fback incorrect cluster 3 | 8 | 0.353 |
| fback incorrect cluster 4 | 5 | 0.398 |

Second, to verify that the resulting networks identified as stable are different than would be obtained by random chance, we shuffled the region labels of all cells before computing the region-wise functional similarity scores and Louvain clustering. The resulting groupings were visually much more numerous and smaller than in the unshuffled data (Fig. S10), and their stability scores were well below the stability score threshold (Table 4).

**Table 4.**
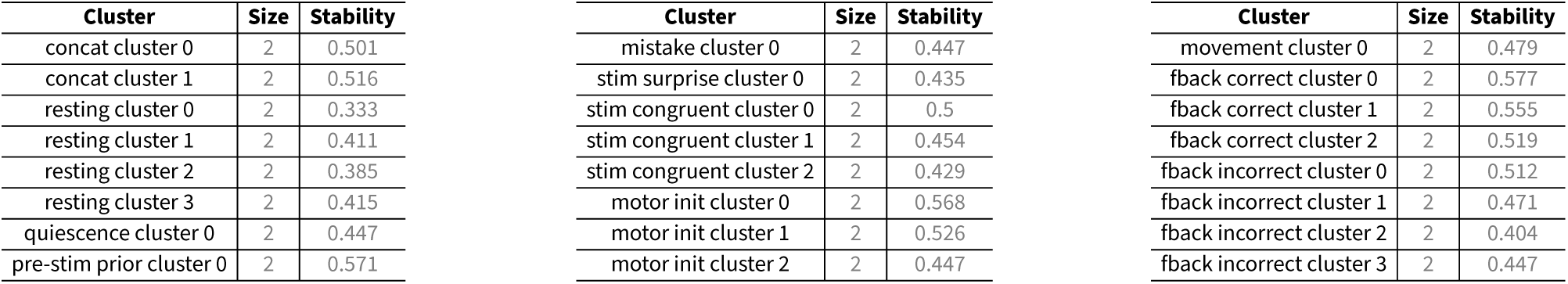
Stability scores for shuffled control clusters. Shuffled clusters generated by shuffling region labels of cells before calculating regional similarity scores and identifying groupings with Louvain algorithm. None of these shuffled clusters show stability scores *>* 0.57 and size ≥ 6 that pass the stability criteria.

| Cluster | Size | Stability |
| --- | --- | --- |
| concat cluster 0 | 2 | 0.501 |
| concat cluster 1 | 2 | 0.516 |
| resting cluster 0 | 2 | 0.333 |
| resting cluster 1 | 2 | 0.411 |
| resting cluster 2 | 2 | 0.385 |
| resting cluster 3 | 2 | 0.415 |
| quiescence cluster 0 | 2 | 0.447 |
| pre-stim prior cluster 0 | 2 | 0.571 |
| mistake cluster 0 | 2 | 0.447 |
| stim surprise cluster 0 | 2 | 0.435 |
| stim congruent cluster 0 | 2 | 0.5 |
| stim congruent cluster 1 | 2 | 0.454 |
| stim congruent cluster 2 | 2 | 0.429 |
| motor init cluster 0 | 2 | 0.568 |
| motor init cluster 1 | 2 | 0.526 |
| motor init cluster 2 | 2 | 0.447 |
| movement cluster 0 | 2 | 0.479 |
| fback correct cluster 0 | 2 | 0.577 |
| fback correct cluster 1 | 2 | 0.555 |
| fback correct cluster 2 | 2 | 0.519 |
| fback incorrect cluster 0 | 2 | 0.512 |
| fback incorrect cluster 1 | 2 | 0.471 |
| fback incorrect cluster 2 | 2 | 0.404 |
| fback incorrect cluster 3 | 2 | 0.447 |

Our third control was based on the statistical significance of the functional responses of the identified networks relative to shuffles. We computed each network’s PETH as a weighted (by region participation scores) average of the PETHs of the member regions, Fig. S9b-e (line plots), and compared these to PETHs generated by randomly sampling the same number of regions from all recorded Beryl regions in the brain (gray lines in the line plots are the shuffled controls). A network was deemed significant in its functional properties if its PETH was significantly different from the random network PETHs for the majority of time bins (≥ 60%).

We find that within each condition, brain states typically form a few non-orthogonal but distinct and regionally distributed similarity subnetworks, suggesting distinct processing modes, Fig. S9b-e. The regional composition of identified subnetworks varies substantially from one task phase and condition to the next, meaning that functional groupings are transient and rapidly shift across time and condition in the task, Fig. S9b-e (Swanson maps, right panels; darker colors indicate higher participation score). For the full concatenated PETHs (“concat”, Fig. S9b), the three dominant subnetworks consist of neocortex-hippocampus, striatum-hypothalamus-midbrain–hindbrain, and cerebellum networks, and while in the congruent stimulus phase (Fig. S9c), the networks collapse to a dominant one, integrating the striatum-hypothalamus-midbrain-hindbrain network with some hippocampal and somato-sensory regions. Movement initiation and movement, adjacent task phases, involve quite different subnetworks, reflecting dynamic reconfiguration (Fig. S9d): the movement initiation period exhibits three distinct subnetworks, one primarily neocortical engaging early sensory processing, somatosensory, and prefrontal areas, but also interacting with lateral hypothalamus, inferior colliculus, and cerebellum; the second is predominantly hippocampal-thalamic-dorsal striatum (CP) with somatosensory-secondary motor cortex engaged with midbrain and hindbrain; and the third is predominantly cerebellar with engagement of superior colliculus. Brain states in the movement period collapse into two statistically robust subnetworks, one (network 2) largely similar to the first movement initiation subnetwork (network 0), and the other a mixture of hippocampus, high-level association areas (RSP), midbrain (PAG), and hypothalamus. Finally, the incorrect feedback phase shows one subnetwork (Fig. S9e) that is similar to the motor-init network 2, presumably reflecting movement-related computations leaking through from the movement phase immediately earlier.

One consistent pattern is that the within-condition subnetworks in every condition recapitulate the green-pink or cortical-subcortical response divide that we saw earlier at the single-cell level. This is evident in the graphs of Fig. S9, as well as when we form a similarity matrix of the region membership of subnetworks from all the task conditions, Fig. S9f. The subnetwork regional similarity matrix has a two-block structure, each with a representative subnetwork from most trial conditions. The upper block corresponds to primarily cortical processing networks, and the lower one to primarily subcortical processing networks. This cortical-subcortical separation is also supported by previous tract-tracing connectomics studies in the mouse brain, showing dense intra-cortical communities are largely segregated from dominant subcortical (thalamic, striatal, hypothalamic, brainstem) communities, with specific hub regions bridging the two compartments ( ^(81;82)^). Finally, we compare these functional brain networks to established fMRI-based networks from the mouse brain (including default mode^(10;48;42;41;45)^, limbic/hippocampal^(43;46;42;47)^, salience^(40;42;45)^, sensorimotor^(46;44;41;42)^, and associated cortical networks^(48;45;42;41)^; see Table 2 for datasets included) to determine whether there is a correspondence, Fig. S9g. The mouse fMRI sensory mode network (smn, Fig. S9g, last row) reassuringly overlaps with a subnetwork from the congruent stimulus conditions, but also with the more-cortical subnetworks from each of the movement initiation and movement conditions. However, beyond this similarity, we report few meaningful relationships between fMRI-defined subnetworks and our high spatio-temporal resolution brain networks. One would expect the resting or quiescent state network would correspond to the fMRI default mode networks (Fig. S9g, bottom row); however none of the quiescent / resting networks are reliable enough to survive the significance tests. The difference of recording conditions (resting state fMRI vs IBL task), the quantified dominance of subcortical processing in the brain throughout the course of the IBL task, the sparsity of fMRI maps in the mapping to Beryl regions even for cortical structures, and the lack of deep subcortical (e.g. midbrain, hindbrain, cerebellum) inclusion in the fMRI network studies are the likely causes of the discrepancy. We also note that our analyses do not measure trial-by-trial or moment-by-moment correlated fluctuations and therefore cannot establish functional connectivity or inter-regional coupling. The observed’reconfiguration’ refers to changes in functional similarity structure across task phases, not direct evidence for changing inter-regional communication.

Together, these results show in a spatiotemporally precise and brain-wide way that all brain areas participate in all functions, and that the brain forms emergent collective dynamical coalitions that rapidly reconfigure between task phases to enable accurate perceptual decision-making.

### Method: functional distance metric for brain region pairs

We measure the functional distance between brain regions as follows. For a given region, the 2D umap embedding of all cells in this region (across recordings) is rasterized into a 256 x 256 pixel grid, with the cell density per pixel, then smoothed by applying a 2D Gaussian kernel with standard deviation of 10 pixels in each dimension and normalized such that the maximum grayscale per pixel is set to 1 and the minimum to zero. Given two such images for two regions, the images are flattened into vectors **v_l_**, **v_2_**and the cosine similarity of the two vectors is defined as the functional distance between regions:

**Figure S9.**
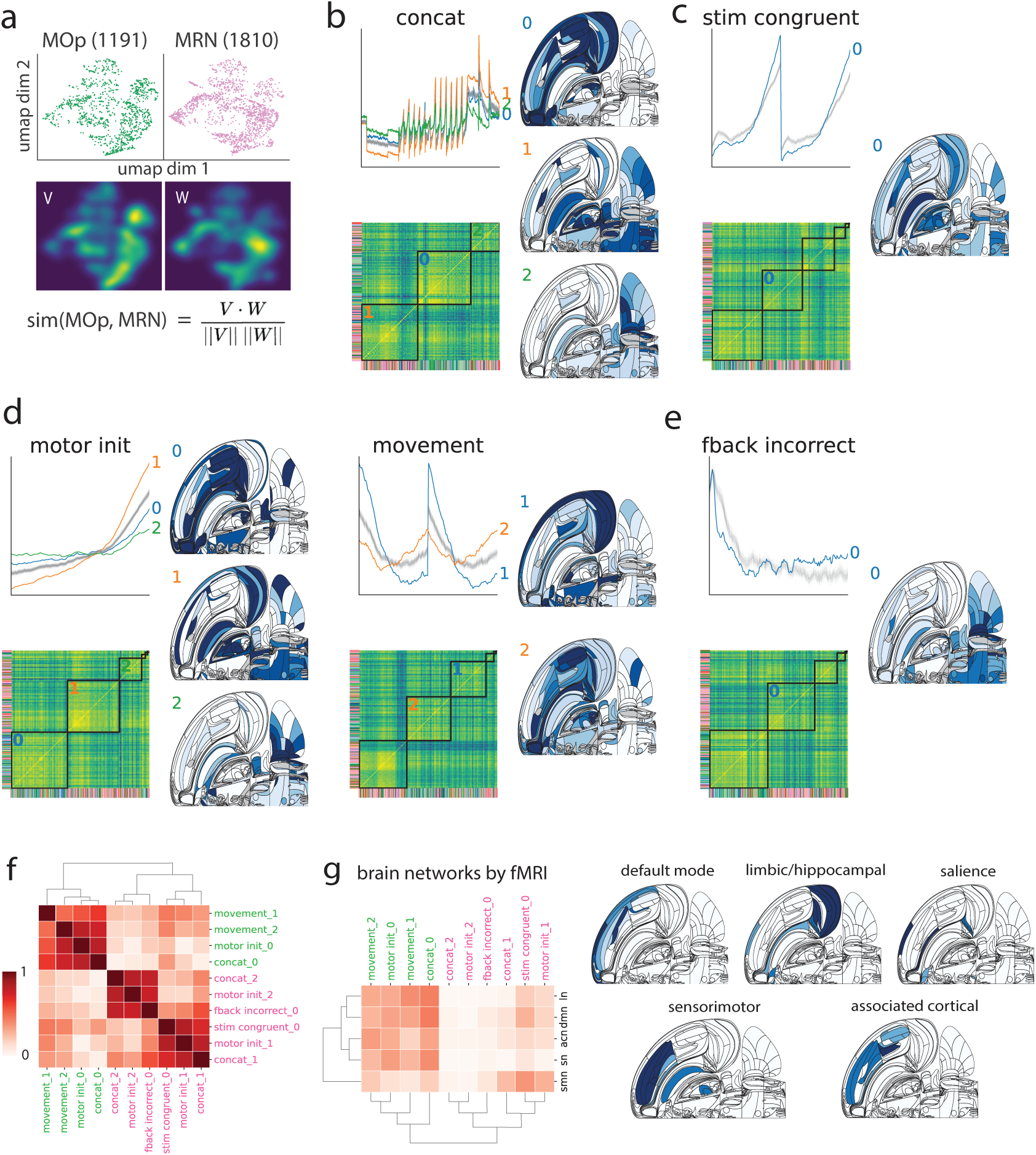
Dynamically reconfiguring networks across different phases of the task. **a** To compare functional responses between pairs of regions, we calculate a cosine similarity score of two regions’ cellular responses from a smoothed 2D embedding for the relevant task phase (here, shown for the concat condition which includes all task phases). **b-e** The regional similarity matrices are ordered by the Louvain community detection algorithm to identify groups of brain regions (networks) for various subsets of PETHs (task phases) shown in panels (b) through (e). The algorithm consistently shows a cortical-subcortical separation in two (or three) major networks across phases of the task, as evident in the Swanson maps (some clusters mostly subcortical, and the others mostly cortical + hippocampus) which show regional membership in these clusters (darker color indicates stronger membership from split-trial cross-validation). We also show line plots of the average PETHs for each network as compared to shuffled controls (gray), and the networks are only included in the current figure if they pass all three rounds of control analyses as described in the Methods. **f** Summary table showing cosine similarity of Swanson maps between pairs of our defined networks, having two clear groups that correspond to the cortical-subcortical separation. **g** Cosine similarity of Swanson maps between our defined networks and canonical fMRI networks in the mouse brain (Swanson maps shown on the right), showing low similarity throughout. We combined multiple studies (see Table 2 for details) for each mouse fMRI network, mapped the results of these studies onto Beryl regions, and colors in the Swanson maps indicate how consistently we find the regions being identified in a network, with darker color being more consistent.

### Method: Louvain clustering

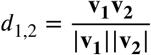

We use sknetwork.clustering.Louvain implemented in python to identify regional networks in the functional similarity matrices of different trial phases. We use the following parameters in the presented analysis: resolution=1.01, random_state=0, after sweeping through parameters and other clustering algorithms that result in less reproducible networks with lower adjusted random index (sklearn.metrics.adjusted_rand_score) and adjusted mutual information (sklearn.metrics.adjusted_mutual_i

### Method: statistical significance of networks

To assess which of the identified networks were reliable and meaningful, we applied three types of control analyses.

First, we identified which networks had a high’network stability’ score through split-trial comparisons, as follows. We split all trials of a given type into two groups to obtain two independent feature vectors for each cell. Within each split, we re-computed similarity matrices and Louvain clusters (communities). We repeated this process four times to generate a total of eight sets of leave-half-out communities for each condition (each cross-validation set based on a random subset of 1∕2 the total trials). We compared region participation across the eight sets of communities to define a network stability score, as follows: All of the communities (for a given condition) in all the eight splits were assigned binary region participation vectors (of length equaling the number of Beryl regions), with 1 indicating a region’s membership in the community and 0 otherwise, and we used these to create a regional membership similarity matrix across all clusters in all splits for the condition (Fig. S10), sorting the matrix elements by similarity to display the communities (SI Fig. S10, column 1). We only look at communities with sizes *>* 6 when sorting the similarity matrix, meaning these communities are identified in at least 6 out of the 8 leave-half-out trial splits analysis. The network stability score of each community is defined as the average value (of the regional membership similarity score) within the community (Table 3): conceptually, if the same areas always appear together in one community across splits, the stability score would be 1; otherwise, it would take some value in the interval [0, 1]. Communities with a stability score *<* 0.57 (the average stability score across communities and across trial phases) were deemed statistically unstable and not numbered or plotted as Swanson maps in the main or supplemental figures.

Second, to verify that the resulting stable identified networks are different than would be obtained by random chance, we shuffled the region labels of all cells before computing the region-wise functional similarity scores and Louvain clustering. The resulting groupings were visually much more numerous and smaller than in the unshuffled data (Fig. S10). We quantitatively compared the shuffled data networks with the identified networks by computing the stability scores for the shuffled data networks. Most of the shuffled networks have grouping size stability scores below the criteria of stability score *>* 0.57, and when comparing them across cross-validation splits, none of them show communities of size ≥ 6(Table 4).

Our third control was based on the statistical significance of the functional responses of the identified networks. The goal is to ensure the networks are a result of distinct and related responses of brain regions within a community rather than community-independent regional response variations. For each community, we computed its cluster PETH by a weighted average of the PETHs of the member regions (weighted by the participation scores of the regions, as defined in the next section in Methods), Fig. S9b-e (line plot in each panel). We compared the community PETH with 1000 control PETHs generated by randomly sampling the same number of regions as in the community from across the set of all recorded Beryl regions in the brain, and computing their average PETHs weighted by the same set of regional participation scores (randomly assigned to the random set of regions), Fig S10. At each time bin, two p-values are defined by the fraction of times the shuffled controls are larger or smaller than the true PETH, and the true PETH is significantly different from the controls if either of the p-values are smaller than 0.01. A community was deemed significant only if the resulting network PETHs were significantly different from the shuffled controls for the majority of the analysis time window.

Only networks that are deemed significant by the intersection of the three conditions above are numbered in Fig. S9 of the main text. In Swanson maps, we show the membership of regions in each significant network, weighted by a network participation score.

### Method: regional participation score of networks

We define a region’s participation score within a given network as the fraction of times the region is identified within the network across the eight-fold split-trial controls. If a region is identified in a given network in all eight splits, its participation score in the network is 1, and if identified in none, its score is 0. For each network that passed the statistical significance tests, we show the Swanson maps of them in the results (Fig.S9) by coloring the regions with the regional participation score.

For each of the mouse fMRI networks, we combined results from multiple studies as shown in Table 2, mapped their included areas onto Beryl regions, and calculated the regional participation score across the studies we included. We show the Swanson maps of each network colored by the regional participation scores in Fig.S9g.

### Cortical–subcortical separation in functional composition space

To quantify the cortical–subcortical distinction in functional cell-type composition, we computed for each Beryl region containing at least 20 neurons a 25-dimensional composition vector **p***_r_*, where *p_r,k_* is the fraction of neurons in region *r* assigned to functional cluster *k* (*K* = 25 k-means clusters). Regions were labelled cortical (Cosmos groups: Isocortex, OLF, HPF, CTXsp; 68 regions) or subcortical (CNU, TH, HY, MB, HB, CB; 130 regions); regions mapped to ‘void’ or ‘root’ were excluded, yielding 198 regions in total.

Three complementary metrics were applied. First, a **weighted Fisher criterion**

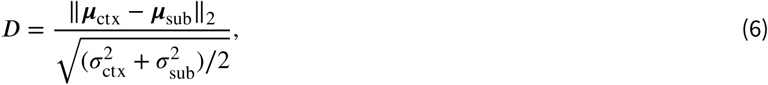

where ***µ*** and *o-*^2^ are the cell-count-weighted centroid and mean squared deviation of regional composition vectors within each group. Weighting by cell count means the result reflects neuron-weighted regional composition rather than treating each region equally; this is the primary analysis because regions with few neurons contribute unreliable composition estimates. Second, **PERMANOVA** (^83^) was applied to the matrix of pairwise Euclidean distances between composition vectors (unweighted), with cortical/subcortical as the grouping factor (scikit-bio, 1000 permutations). Effect size *R*^2^ was derived from the pseudo-*F* statistic as *R*^2^ = *F* ∕(*F* + *n* − 2), where *n* is the number of regions. Third, **PCA** was performed on the composition matrix (rows weighted by ^√^*n_r_* prior to fitting) and the first two principal components were visualised. For all permutation tests, empirical *p*-values use the +1 correction: *p* = (1 + #{*T*_perm_ ≥ *T*_obs_})∕(1 + *N*_perm_), giving a minimum resolvable *p* of 0.001 for 1000 permutations.

**Table 5.** Visual cortical regions included in the dataset. For each Allen visual cortical area, the table reports the region acronym, full region name, number of recordings, total number of recorded neurons, and number of neurons passing the “good neuron” quality criterion. Find this information for all regions at (^75^).

| Acronym | Region name | # recordings | # neurons | # good neurons |
| --- | --- | --- | --- | --- |
| VISal | Anterolateral visual area | 5 | 697 | 95 |
| VISam | Anteromedial visual area | 26 | 2581 | 374 |
| VISl | Lateral visual area | 14 | 1559 | 151 |
| VISp | Primary visual area | 65 | 8559 | 880 |
| VISpl | Posterolateral visual area | 14 | 523 | 95 |
| VISpm | Posteromedial visual area | 22 | 1995 | 241 |
| VISli | Laterointermediate area | 9 | 656 | 83 |
| VISpor | Postrhinal area | 8 | 889 | 99 |

**Figure S10.**
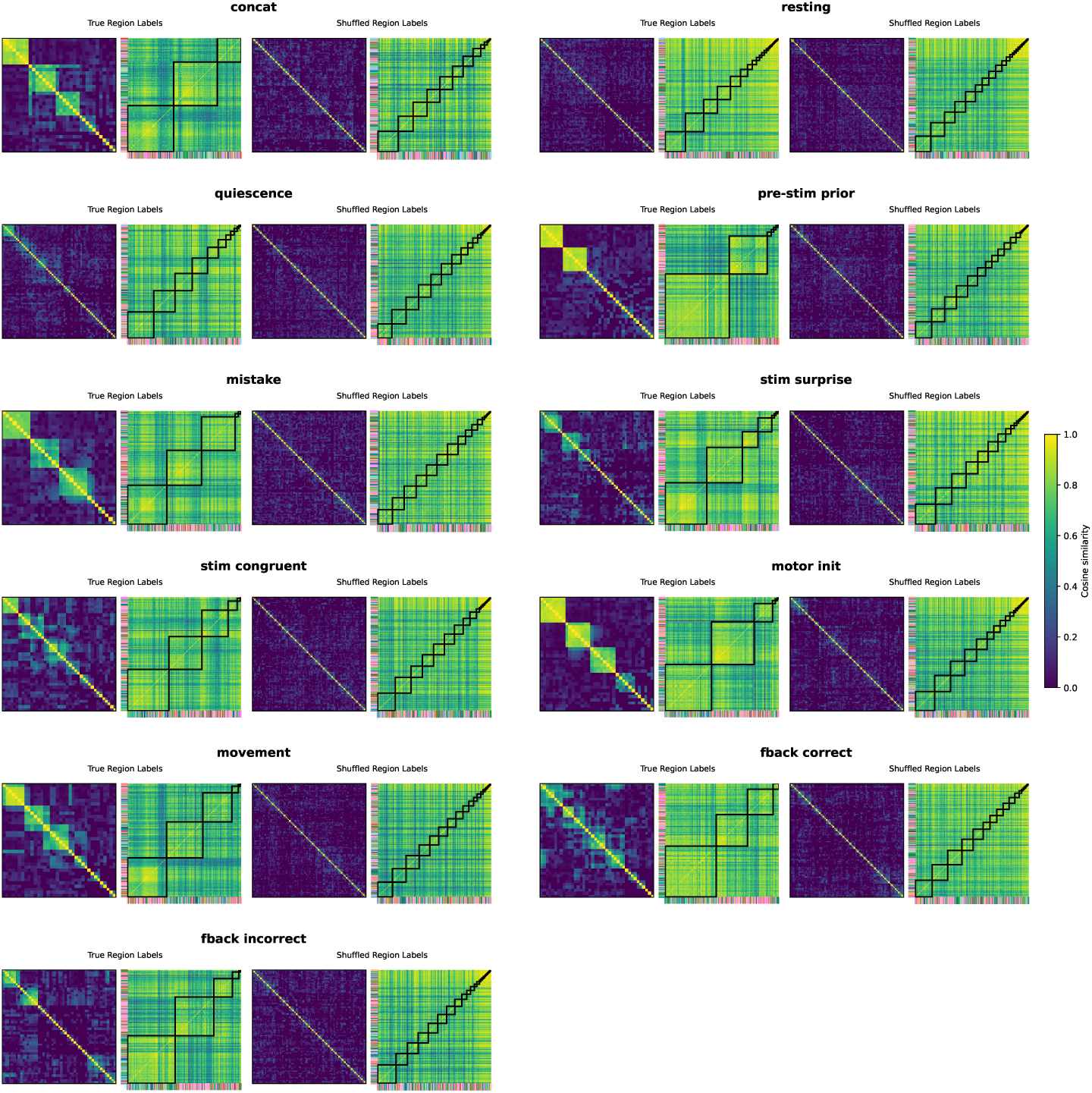
Shuffled controls for Louvain community detection. Created by randomly shuffling region labels of all cells before computing the functional similarity scores between regions and running Louvain algorithm to identify clusters. For each trial phase, we show the resulted clusters from the true region labels (2nd column of each panel) and those from shuffled region labels (4th column of each panel), and compare their reliability by computing cosine similarity between all sets of clusters generated from the 8-fold split-trial cross-validation process (1st and 3rd columns of each panel; similarity score matrices ordered by hierarchical clustering). Using the true region labels, Louvain algorithm detects some reliable clusters that are consistent across different splits of trials (relatively big clusters in similarity matrices, 1st column of each panel; true for most trial phases except resting), whereas controls using the shuffled region labels always result in inconsistent clusters (3rd column of each panel).

**Figure S11.**
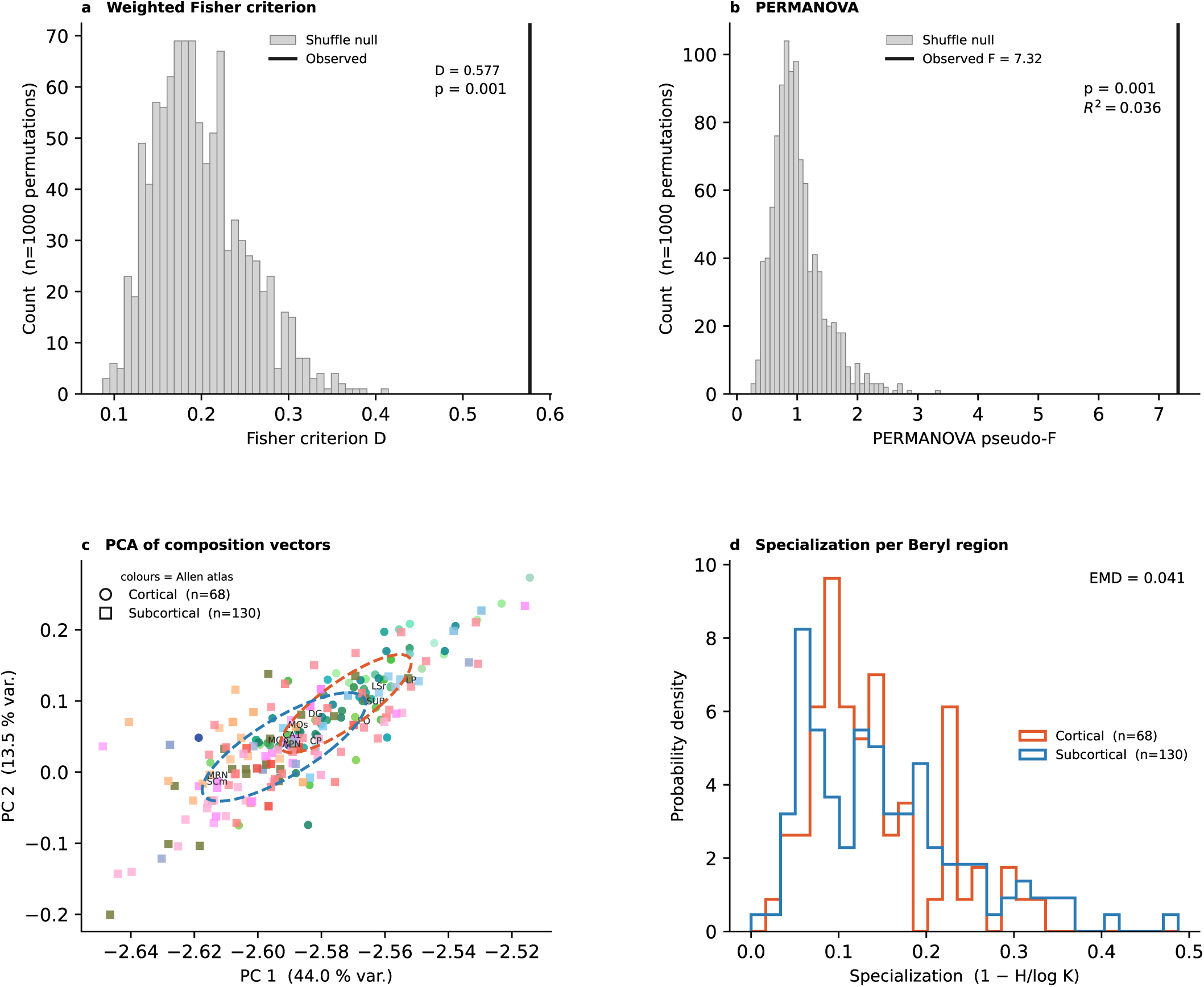
Cortical–subcortical separation in functional-cluster composition space. For each of 198 Beryl regions (≥20 neurons), we computed the fraction of neurons assigned to each of 25 cross-validated k-means functional clusters. **a** Weighted Fisher criterion comparing cortical and subcortical composition vectors. The observed separation (*D* = 0.577; vertical line) exceeded all 1000 label-shuffle permutations (*p* = 0.001). **b** PERMANOVA on pairwise Euclidean distances between regional composition vectors confirmed a significant but small group effect (pseudo-*F* = 7.32, *R*^2^ = 0.036, *p* = 0.001). **c** PCA of composition vectors (rows weighted by ^√^*n*_cells_; PC1: 44.0%, PC2: 13.5%). Points are Beryl regions coloured by Allen atlas colour; circles denote cortical and squares subcortical regions; dashed ellipses show 1 s.d. within-group scatter. Cortical/subcortical separation is significant in the full composition space but modest relative to the dominant shared axes of variation, consistent with broad anatomical mixing of functional response types and residual differences in their proportions. **d** Specialization score (1 − *H*(**p**)∕ log *K*) per Beryl region, separately for cortical (*n* = 68) and subcortical (*n* = 130) regions. Normalised EMD = 0.041. Compare with Fig.S6.

**Figure S12.**
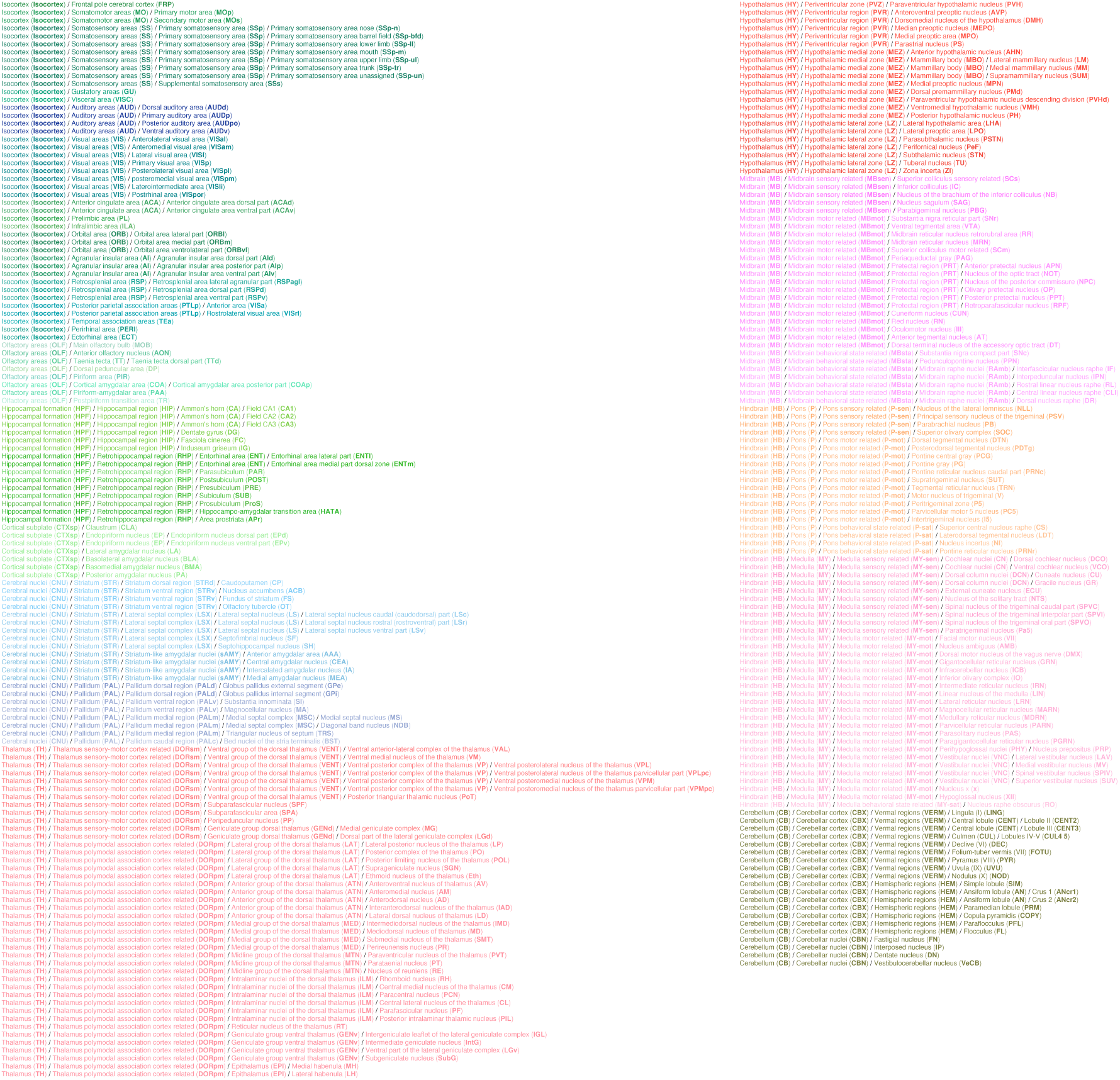
Anatomical hierarchy for all used regions. For each of the used regions, at “Beryl” hierarchical level, the parent structures are shown with full names and acronyms, up to “Cosmos” hierarchical level. Find the full anatomical hierarchy tree at the Allen adult mouse brain atlas.

